# CD33 Epitope Editing Unlocks UM171-Expanded Cord Blood Grafts for AML Immunotherapy

**DOI:** 10.1101/2025.05.07.652621

**Authors:** Bernhard Lehnertz, Maéva Langouët, Sophie Corneau, Tara MacRae, Jean-Francois Spinella, Nadine Mayotte, Maju Joe, Edward N. Schmidt, Susan Moore, Maria Florencia Tellechea, Isabel Boivin, Loïc Papineau, Shanti Rojas-Sutterlin, Margaux Tual, Camille Jimenez Cortes, Jalila Chagraoui, Elisa Tomellini, Virginie Desse, Etienne Gagnon, Matthew S. Macauley, Guy Sauvageau

## Abstract

Immunotherapies in acute myeloid leukemia (AML) are limited by shared antigen expression between leukemic and healthy hematopoietic cells, leading to on-target toxicity. Here we developed a clinically scalable strategy to engineer cord blood (CB)-derived hematopoietic stem and progenitor cell (HSPC) grafts resistant to CD33-directed therapies. Leveraging UM171-mediated expansion and adenine base editing, we precisely disrupted a critical epitope in CD33 required for gemtuzumab ozogamicin (GO) binding, centered on phenylalanine 21, while preserving CD33 expression and its sialic acid binding function. *Ex vivo* edited HSPCs maintained robust multilineage engraftment, T-cell output, and conferred protection from GO-induced myelotoxicity in xenograft models, without impairing anti-leukemic efficacy. Editing was efficient across multiple donors, enriched in primitive subsets, and exhibited minimal off-target activity by ultra-deep exome sequencing. Our work establishes base editor-driven epitope engineering as an improved approach to CD33-targeted immunotherapy-compatible HSPC grafts, enabling safe integration of currently available agents into post-transplant care.

## Introduction

Cord blood (CB) derived hematopoietic stem cell (HSC) grafts offer unique immunologic advantages for allogeneic transplantation, including low rates of graft-versus-host disease^1^ and one of the most potent graft-versus-leukemia activities^2,3^. However, their clinical use has historically been limited by low stem cell dose and delayed engraftment^4^. The small molecule UM171 enables robust *ex vivo* expansion and rejuvenation of CB-derived CD34⁺ cells by promoting degradation of the CoREST1 complex and c-MYC, enhancing self-renewal and engraftment potential^5–7^ overcoming this limitation. Clinically, UM171-expanded grafts lead to reduced non-relapse mortality through accelerated neutrophil recovery^8^ and improved donor-recipient HLA matching^9^. UM171 grafts also maintain the benefits of CB transplants, namely low relapse and cGVHD^10^ possibly because they procure robust NK and CD4 T-cell reconstitution^11^. These benefits have translated into promising outcomes in high-risk AML populations, including patients with TP53-mutant disease or prior transplantation, where UM171 expansion has been associated with 2-year survival rates exceeding 60% (unpublished, presented at ASH 2023, 2024). Nonetheless, disease relapse remains a major challenge, underscoring the need for safe and effective post-transplant immunotherapeutic strategies.

Targeted immunotherapies such as antibody-drug conjugates (ADCs), bispecific T-cell engagers (BiTEs), and CAR T cells are being actively explored in AML, but their clinical use is often limited by antigen overlap between leukemic and normal hematopoietic cells. CD33, the target of the FDA-approved ADC gemtuzumab ozogamicin (GO), is highly expressed on AML blasts but also on healthy hematopoietic stem and progenitor cells (HSPCs) and mature myeloid cells, resulting in dose-limiting myelotoxicity^12,13^. This lack of antigen selectivity similarly hampers the efficacy and safety of CD33-directed BiTEs (e.g., AMG 330^14,15^) and CAR T cells^16,17^. One possible solution is the genetic engineering of HSC grafts to eliminate CD33 expression^18–20^ or delete its second exon^21^.

In this study, we build on the UM171 CB HSC expansion platform to develop a clinically scalable, base editing-driven approach for engineering CD34⁺ HSPC grafts resistant to CD33-directed immunotherapies. We identify a precise epitope within CD33 required for GO binding and demonstrate that base editing can selectively disrupt this epitope while preserving CD33 expression and graft function. Our work establishes a high-fidelity and integrated method for UM171-expanding and protecting hematopoietic grafts from immunotherapy-induced toxicity, enabling safer and more efficacious post-transplant application of targeted agents in AML.

## Results

### Establishing Optimal HSPC Graft Engineering Parameters

To address limitations of immune target selectivity in AML, we explored best strategies for target epitope ablation in HSCs, enabling immunotherapeutic agents to specifically target leukemia cells while sparing HSC graft-derived cells. In cases where surface molecules are functionally dispensable, as was suggested for CD33^22^, complete gene ablation offers a promising solution^18,19,23,20^. To this end, we benchmarked common CRISPR-based approaches, specifically Cas9 and cytosine base-editor (CBE)-mediated mutagenesis, for complete CD33 ablation. In the context of UM171 supported *ex vivo* expansion of cord blood derived CD34+ HSPCs (**Fig. 1A**), these methods consistently achieved high CD33 knockout efficiency (>90% of cells, **Fig. 1B**) in the context of optimized Cas9/base editor mRNA architecture, chemistry, sgRNA design, and electroporation parameters^24^. Despite achieving high knockout efficiency, Cas9-mediated editing, dependent on DNA double-strand breaks and error-prone repair, significantly impaired CD34^+^ cell expansion (**Fig. 1C**) and graft functionality (**Fig. 1D**). Strikingly, this manifested as a 22-fold and 17-fold reduction in myeloid and B-lymphoid engraftment, respectively (**Fig. 1E**, green versus blue top two facet rows, week 21 post-transplant). Additionally, we observed a near-complete loss of T-cell output (**Fig. 1E**, green versus blue in bottom facet row, week 21 post-transplant, **suppl. Table 1**). In contrast, cytosine base editor (CBE)-mediated nonsense mutation^25^ of tryptophan 60 (W60X), which avoids DNA double-strand breakage and instead converts guide RNA-targeted cytidines to uridine/thymidine via deamination^26^, had little impact on graft expansion (**Fig. 1C**, purple versus blue), and no significant impact on graft activity in NSG mice (**Fig. 1D**). These results suggested that CBE-mediated CD33 ablation preserves the full range of benefits offered by UM171-expanded cord blood HSPCs in xenografted NSG mice, including high and uniform engraftment levels (**Fig. 1D**, unedited and edited +UM171 versus unedited-UM171), lympho-myeloid lineage balance characteristic of young HSCs, and robust T-cell potential essential for graft-versus-leukemia activity. Based on these experiments, we concluded that base editing represents a superior *ex vivo* gene editing strategy for HSC graft engineering.

**Figure 1:**
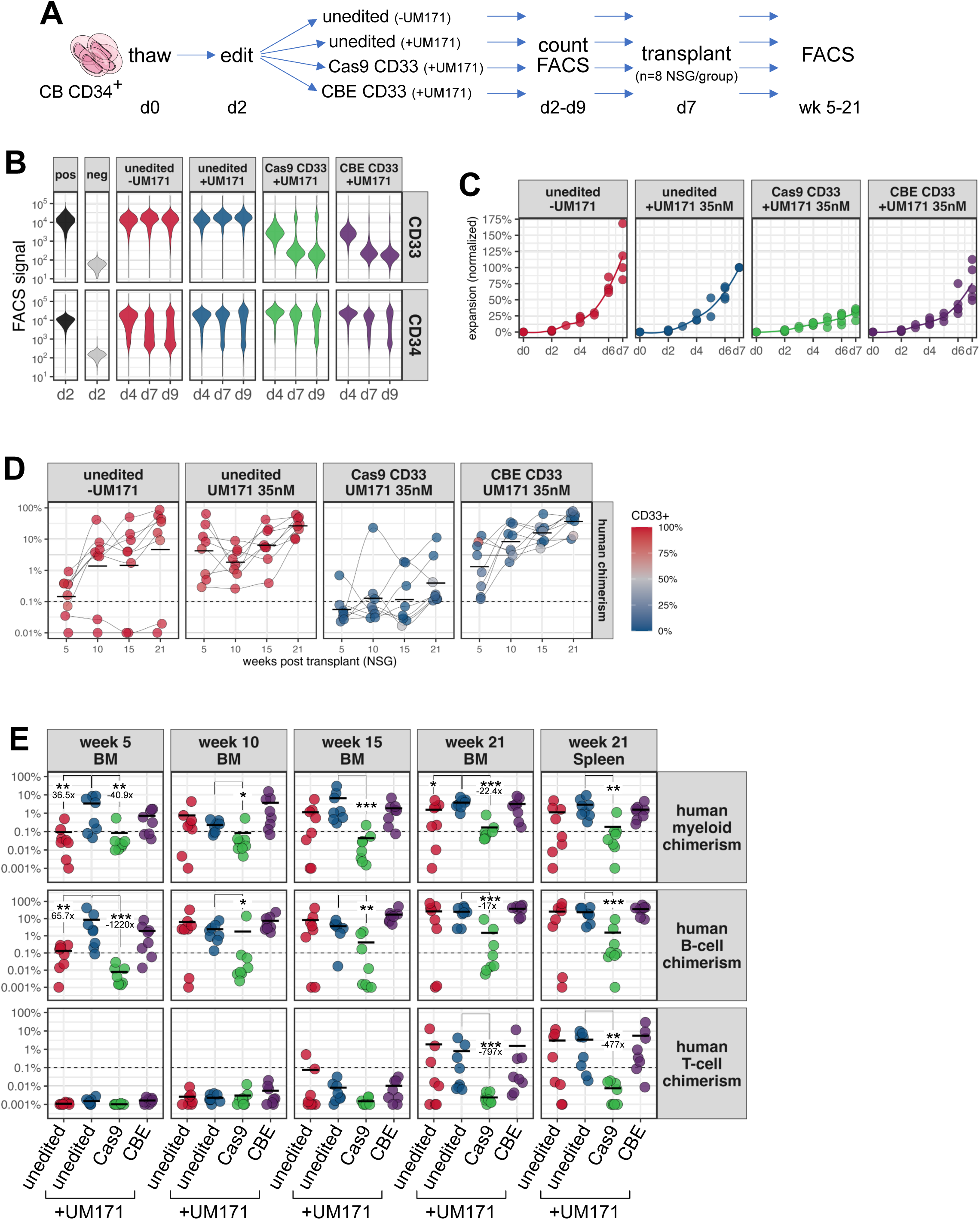
Benchmarking of CD33 Ablation Strategies in UM171-Expanded Cord Blood CD34+ Cells. A: CD34+ CB cells were expanded ± UM171 (35nM), and CD33 and CD34 levels were measured by FACS at indicated timepoints. Cas9 or CBE mRNAs, in combination with a CD33 exon 2 targeting sgRNA were electroporated at d2 of culture to introduce frameshift or W60X missense mutations, respectively. **B:** CD33 ablation was very efficient in both approaches, indicated by progressive loss of surface expression. CD34 expression was preserved in the presence of UM171, with no overt differences between unedited and edited conditions. One representative of five experiments shown. **C:** Total cell expansion of CD34+ CB cells expanded ± UM171 is plotted over 7 days. Gene editing (mRNA + sgRNA electroporation) was done at d2. Combined and normalized data (rel. to unedited +UM171 d7) from 4 independent experiments. A significant impact of Cas9 but not CBE on overall cell expansion was observed. **D:** After 7d of *ex vivo* expansion, unedited CD34+ cells (+/-UM171), Cas9 or CBE edited cells were transplanted at day 0 equivalent doses (progeny of 7K cells) into NSG hosts and monitored over time. Efficient CD33 ablation was observed in both edited conditions, color-scale indicates CD33 expression in graft-derived myeloid cells (CD13+ or CD14+). A significant reduction of human engraftment was observed in Cas9 edited grafts, while base-edited grafts were comparable with unedited UM171-expanded grafts. Crossbars indicate mean, connecting lines indicate different mice across timepoints. **E:** Human lineage positive cells were determined and plotted by timepoint. Statistically significant differences were calculated by Wilcoxon tests relative to unedited +UM171 at each timepoint and tissue (BM = bone marrow). Significant Benjamini-Hochberg adjusted p-values are indicated as *<0.05, **<0.01 and ***<0.001. No significant differences between unedited and base-edited UM171-expanded grafts were detected. Human myeloid and B-cell potential is greatly impaired and T-cell potential is essentially lost in Cas9 edited grafts. Crossbars indicate mean. Full statistics summary is provided in supplemental table 1. One representative of five total experiments is shown.

### CD33 Epitope Engineering to Evade GO On-Target Toxicity

Given the demonstrated feasibility of UM171-expansion and base-editing in HSC graft engineering, we next sought to introduce targeted missense mutations in CD33 to eliminate cross-reactivity with the therapeutic CD33 antibody clone P67 used in GO/Mylotarg, while preserving CD33 expression and function. Since CD33’s biological role appears dispensable for HSCT but has been shown to suppresses innate immune cell reactivity^27^, we reasoned that this approach may present important safety benefits over complete gene inactivation. Similar approaches have recently been reported for AML immune targets with essential functions in stem and progenitor cells, such as KIT, FLT3^28^, CD123^29^ and CD45^30,31^.

To identify base-editable epitopes in CD33, we first designed a high-density lentiviral sgRNA library to target most amino acids in the extracellular V-set domain of CD33 encoded in exon 2 (**Fig 2A**, **suppl. Table 2**), since binding of the therapeutic CD33 P67 and diagnostic WM53 antibodies have previously been mapped to this region^32^. We then transduced CD33-expressing AML5 cells at a low infection-rate (∼20%), followed by drug selection and electroporation with mRNAs encoding either PAMless^33^ adenine (SpRY ABE8e^34^) or cytosine base editors (SpRY TadCBEd^35^). CD33 P67 and WM53 co-staining identified two populations that selectively lost recognition by either one of these antibody clones in ABE but not in CBE-edited conditions (**Fig 2B**, middle panel, red and blue gates and **suppl. Fig 1**). These populations also maintained cross-reactivity with the HIM3-4 CD33 antibody clone (**Fig 2B**, right panel), which was shown to bind outside of the library-targeted region, confirming retained CD33 surface expression.

**Figure 2:**
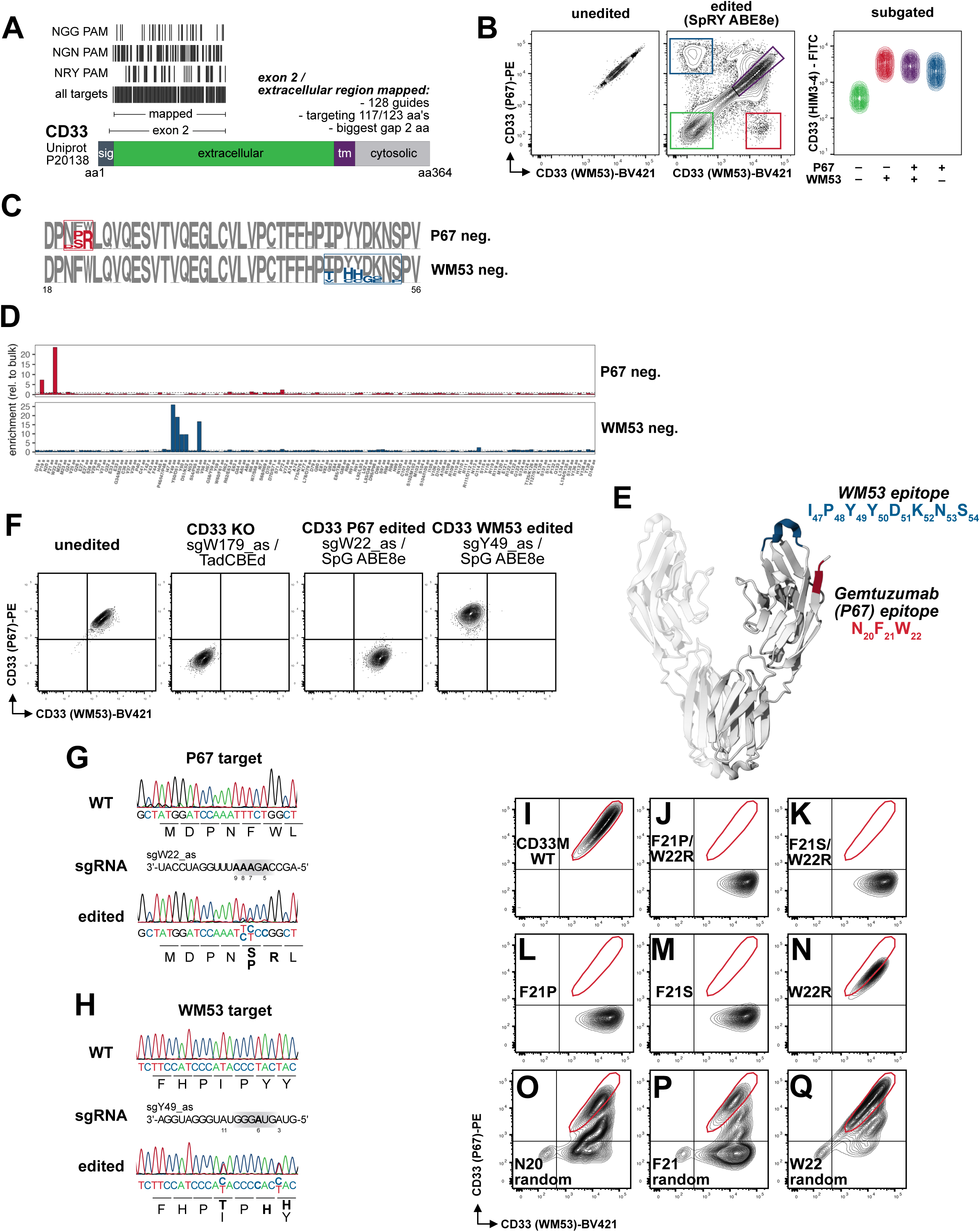
Base Editable CD33 GO Epitope Identification. A: An sgRNA library was designed to densely map the extracellular domain of CD33 encoded in exon 2 as both CD33 antibodies (WM53 and P67) were previously shown to bind this region. The locations of individual sgRNAs falling into three different PAM contexts (NGG, NGN and NRY) are represented as black vertical bars. Library summary including sequences, predicted mutations, PAM contexts etc. is provided in supplemental table 2. Sig: signaling peptide; tm: transmembrane domain, **B:** Puromycin-selected sgRNA library expressing AML5 cells were either left unedited or transfected with a PAMless (SpRY) ABE8e encoding mRNA. Double staining with WM53 and P67 anti-CD33 antibodies revealed presence of single positive populations, carrying missense mutations abrogating binding of either antibody. The population gated in red carried sgRNAs capable of mutating a three amino acid CD33 P67 bound region close to the extracellular N-terminus of CD33, and the population gated in blue identified an eight amino acid CD33 WM53 epitope region, highlighted in **C, D** and **E**. **F:** In combination with SpG ABE8e, single validated sgRNAs were capable of completely blocking recognition by either P67 or WM53 anti-CD33 antibody clones in U937 cells. CD33 P67 (**G**) and WM53 (**H**) de-epitoped and FACS sorted cells were sequenced on CD33 exon 2, revealing minimal missense mutations required for antibody de-epitoping. **I-Q:** CD33M-lentiviral rescue experiments in CD33 KO U937 cells confirmed complete loss of CD33-P67 recognition in the two ABE8e generated double variants (**J** and **K**). Single mutant recue experiments determined that CD33 F21 was the most critical amino acid for recognition by anti CD33 P67 (**L-Q**).

Targeted sequencing of CD33 exon 2 in bulk, CD33 P67 or CD33-WM53 single-positive populations highlighted missense mutations in a three amino acid window close to the extracellular N-terminus of CD33 (N_20_F_21_W_22_) identifying a CD33 P67 epitope (**Fig 2C and D**, red) as previously suggested^36^, and a region of eight amino acids (I_47_P_48_Y_49_Y_50_D_51_K_52_N_53_S_54_) likely recognized by the CD33 WM53 antibody clone (**Fig 2C and D**, blue). These regions locate in relative 3D proximity to each other (**Fig 2E**), explaining the partial binding competition between CD33 P67 and WM53 antibodies^32^. Furthermore, sgRNAs targeting these regions were also enriched in the CD33 P67 and WM53-negative populations, respectively (**Fig. 2D**). We validated these results through electroporation of single synthetic sgRNAs alongside mRNAs encoding either canonical SpCas9, SpG, or SpRY ABE8e, depending on the respective PAM contexts (**suppl. Fig. 2**). These experiments identified the most potent sgRNAs for disrupting the CD33 P67 and WM53 epitopes, designed to target either tryptophan 22 or tyrosine 49 through antisense sgRNAs, respectively (**Fig. 2F and G, suppl. Fig. 2**). In addition to the primary intended missense mutations in these designs, bystander mutations in adjacent codons occurred in both the CD33 P67 and WM53 epitope editing reactions. In CD33 P67-edited cells, the W22R mutation, caused by A-to-G transition in the antisense binding protospacer position 5 (**Fig. 2G**, left), was accompanied by either F21P or F21S mutations caused by complete deamination of protospacer adenine 7 and partial deamination of adenines 8 and 9, respectively. Similarly, in WM53-edited cells, the primary target tyrosine 49 (Y_49_) was invariably converted to histidine, whereas isoleucine 47 was converted to threonine and tyrosine 50 was converted to histidine in about 50% of alleles, respectively (**Fig. 2G**, right).

We next performed a detailed characterization of the CD33 P67 epitope using lentiviral rescue experiments in CD33 KO U937 cells (**Fig. 2I-Q**). Overexpression of exon 2 including (CD33M) variants confirmed that the P21/R22 (**Fig. 2J**) and S21/R22 (**Fig. 2K**) double mutants selectively lost CD33 P67 cross-reactivity (**Fig. 2I**). Interestingly, the F21P (**Fig. 2L**) and F21S (**Fig. 2M**) single mutations also fully disrupted the CD33 P67 epitope, whereas W22R alone only partially attenuated cross-reactivity (**Fig. 2N**). These results demonstrated that ABE8e-mediated CD33 P67 epitope disruption is primarily achieved by mutagenesis of F21.

To further dissect the contribution of individual residues in the N_20_F_21_W_22_ motif, we introduced codon-randomized lentiviral pools for N20, F21, or W22 (**Fig. 2O, P, and Q**). These experiments confirmed that F21 was essential for CD33 P67 antibody binding, as most cells in this codon pool completely lost cross-reactivity (**Fig. 2P**). In contrast, CD33 P67 epitope loss was less frequent and largely incomplete in the N20 (**Fig. 2O**) and W22 (**Fig. 2Q**) randomized pools. Accordingly, narrow-window ABEs^37,38^ failed to generate completely de-epitoped alleles in both CD33 P67 epitope (sgW22_as) and WM53 epitope (sgY49_as) targeted editing reactions (**suppl. Fig. 3**).

### CD33 P67 Epitope Variants Retain Sialic Acid Binding Capacity

As member of the Siglec family, CD33 engages with self-associated sialylated glycan structures on the same (*in cis*) or on opposing cells (*in trans*), resulting in its activation/internalization to dampen immune reactivity towards healthy host cells^39,40^. To evaluate the functional implications of epitope-engineered CD33 variants, we measured their capacity to bind a human CD33-selective modified sialoside ligand^41^ (**Fig. 3A**, CD33L), displayed on Alexa-647 labeled liposomes^42^, mimicking *trans* interactions (**Fig. 3B and C**). CD33 P67-edited U937 cells largely retained the ability to bind this ligand and trigger CD33 internalization, albeit with some attenuation compared to both unedited and WM53 edited cells (**Fig. 3D and E**, red vs blue bars, **suppl. Table 3**). Furthermore, lentiviral rescued CD33 KO U937 cells generally confirmed these results and further highlighted that CD33M P21R22 performed favorably over the CD33M S21R22 variant in these assays (**Fig. 3F and G**, **suppl. Table 3**). Naked liposomes (blue), CD33 KO and the sialic acid binding deficient CD33M R119A mutant^43^ served as controls, demonstrating selectivity. Of note, double mutants in U937 cells were associated by increased surface expression, both in endogenous and lentiviral transgene contexts (**Fig. 3E and G**, blue bars in lower facets).

**Figure 3:**
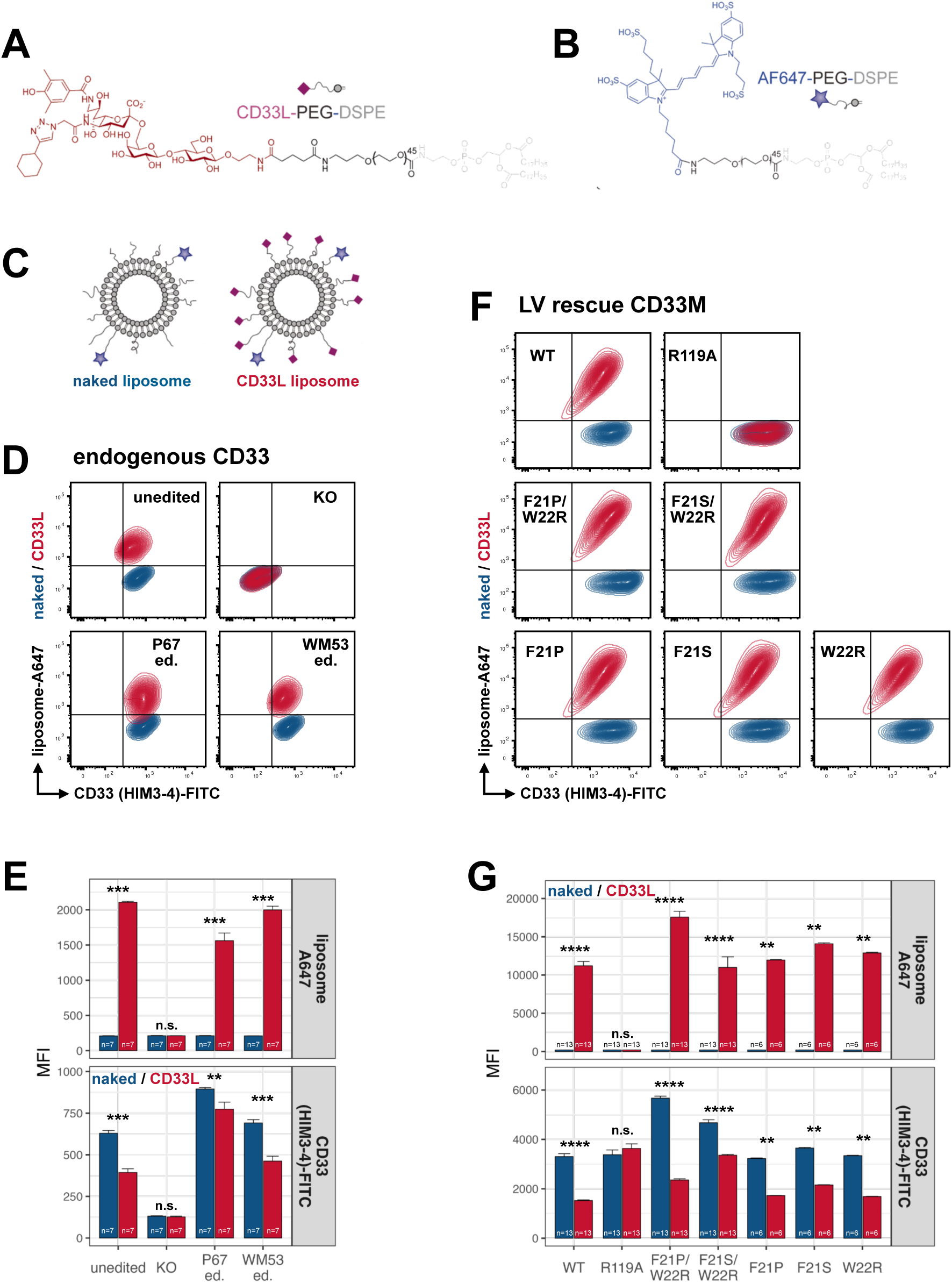
Functional Assessment of CD33 P67 Epitope Variants. A-C: Fluorophore conjugated CD33L liposome formulation for engaging CD33 on cells. Chemical structures of CD33L-PEG-DSPE (**A**), Alexa-647 (A647)-PEG-DSPE (**B**), cartoon diagrams of CD33L and naked liposomes (**C**). **D-G:** Binding studies of naked control (blue) and CD33L conjugated liposomes (red) to WT and CD33 KO control U937 cells (top panels), and CD33 de-epitoped U937 cells. Binding of liposomes to cells and their impact on CD33 surface expression was quantified by flow cytometry (**D** and **E**). Rescued CD33 KO U937 cells were analyzed in the same fashion (**F** and **G**). In contrast to the R119A control mutant, previously identified critical for sialic acid binding, P67 epitope edited CD33 variants retained this ability, albeit with some attenuation. MFI = mean fluorescence intensity. Statistical significance was determined by Wilcoxon-test, p-values are indicated by asterisks: ** indicates p < 0.01, *** indicates p < 0.001, **** indicates p < 0.0001, n.s. indicates p >= 0.05. Bars indicate means with standard error bars. Two independent experiments are summarized in **E** and **G**, respectively. Full statistics summary is provided in supplemental table 3.

### Maintained Graft Activity and Protection Against CD33-Directed Immunotherapy

To validate our CD33 de-epitoping strategy in a UM171-expanded cord blood transplantation setting, we used conditions as in **Fig 1A**. We combined the sgRNA targeting the CD33 codons F_21_ and W_22_ with an NG-PAM compatible adenine base editor (SpG ABE8e^33^) also carrying the Cas9 R691A high-fidelity mutation for reduced off-target activity^44^. This resulted in high efficiency editing with the previously used (1µg mRNA/ 1µg sgRNA per 2e6 cells, 1x) and 2-fold increased (2µg mRNA/ 2µg sgRNA, 2x) dose (**Fig 4A**). We routinely observed an intermediate, likely heterozygous population (**Fig 4B**). Transplantation of unedited versus edited cells into NSG mice indicated high persistence of edited myeloid cells (**Fig 4C**), normal engraftment (**Fig 4D**) and lineage potential (**Fig 4E**) of edited HSPCs with a 1x, and only mildly reduced engraftment of cells edited with a 2x mRNA/sgRNA dose (**Fig 4D**, right panel, **suppl. Table 4**). Importantly, the overall proportion of edited cells in engrafted mice was higher than detected at the time of transplantation, likely due to more pronounced editing efficiency in rare immature cells.

**Figure 4:**
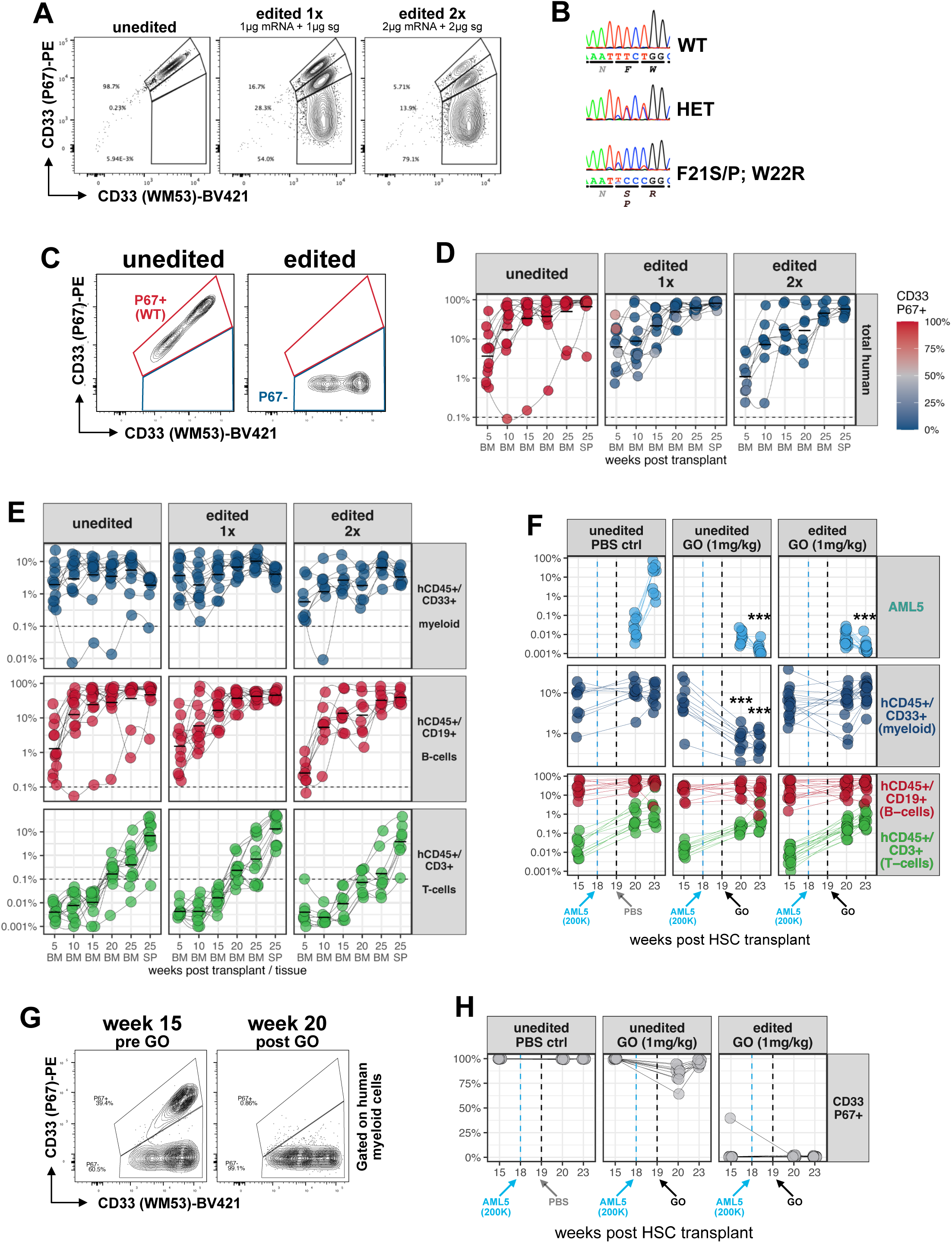
CD33 P67 Epitope Engineering in CD34^+^ HSPCs. A: CB CD34^+^ cells were electroporated at d2 of 7d UM171 expansion culture with an sgW22_as and SpG ABE8e base editor encoding mRNA at two different doses (amounts per 200K cells are shown). FACS analysis (d7) indicated high frequency of editing based on selective loss of P67 binding and sequencing of heterozygous/homozygous mutant populations (**B**). Cells from **A** were transplanted into NSG mice (progeny of 10K CD34+ cells plated at d0). **C:** Example of graft-derived CD33^+^ cells. Normal levels (**D**) and differentiation (**E**) in edited grafts. Color-scale in **D** represents P67 cross-reactivity on graft-derived CD33^+^ cells; colors in **E** represent human B-cells (red), human CD33^+^ (WM53) myeloid cells (blue) and human T-cells (green). Crossbars indicate mean, connecting lines indicate different mice across timepoints. n=12 for unedited and edited (1x) conditions, n=8 for edited (2x). **F:** NSG mice engrafted with either unedited or edited CB CD34^+^ cells received 2*10^5^ AML5 cells 18 weeks post HSC transplant. Control (PBS) or GO treatments were done one week later. Early (week 20) and late (week 23) follow-up bone marrow biopsies indicated significant myelotoxicity (dark blue) in unedited but not in P67 epitope edited cohorts. Anti-leukemic activity of Mylotarg/GO was confirmed (AML5, light blue). ***BH adj. p<0.001 Wilcoxon-test. n=10 for unedited (GO/PBS) cohorts; n=16 for edited GO cohorts (n=9 edited 1x and n=7 edited 2x combined). One representative experiment of two is shown. **G:** Example of a CD33 P67 epitope edited HSC graft recipient with partial persistence of wildtype cells at week 15 post-HSC transplant, which were completely eradicated by Mylotarg/GO by week 20, 7 days post treatment. **H:** Summary of CD33 P67^+^ expression in WM53^+^ myeloid cells. Reduced signal in unedited GO-treated mice was observed due to CD33 internalization. Population percentages and statistics are provided in supplemental table 4.

We next tested the functionality of CD33 P67-edited HSC grafts *in vivo*, in an experimental setup modeling a scenario that presents a severe treatment challenge in the clinic, AML relapse following HSC transplantation. To this end, we inoculated unedited and edited HSC engrafted recipients with CD33-positive AML5 cells, followed by PBS or GO/Mylotarg treatment. Strikingly, edited HSC graft-derived CD33^+^ myeloid cells were significantly protected against Mylotarg/GO (**Fig 4F**, dark blue population, right panel), while unedited counterparts were depleted more than 10-fold for up to four weeks (**Fig 4F**, dark blue population, middle panel, p<0.0001 compared to week 15, **suppl. Table 4**). Importantly, AML5 cells were efficiently cleared (**Fig 4F**, light blue, p<0.0001 compared to PBS controls, **suppl. Table 4**), indicating high on-tumor activity. Further, in a recipient with remaining unedited human cells, GO-treatment depleted those cells with no sign of overall myelosuppression (**Fig 4G** and **H**), demonstrating that even partially edited grafts can achieve functional resistance to GO.

In summary, these results demonstrate that the CD33 P67 epitope engineering strategy achieves highly efficient graft engineering, does not compromise HSC graft expansion or activity following transplantation, and renders graft-derived cells insensitive to even high doses of GO/Mylotarg *in vivo*.

### Reproducibility, Feasibility and Safety Assessment of CD33 P67 HSC Editing

To evaluate clinical feasibility, we aimed to provide evidence of process reproducibility and safety. To this end, CD34⁺ cell preparations from five different cord blood donors were subjected to paired manufacturing runs using unedited and CD33 P67 epitope-edited protocols. Across all runs, the impact of editing on *ex vivo* expansion kinetics was acceptable for sufficient cell yields (**Fig. 5A**). Notably, one experiment (CB 917) showed reduced phenotypic editing efficiency (**Fig. 5B**). Since non-GMP electroporation equipment was used, parameter recording was limited, preventing us from determining if this deviation was technical. The editing rate was reproducibly highest in phenotypically immature subsets (**Fig. 5B**), consistent with extended mRNA stability in these slow-proliferating cells (**suppl. Fig. 4**). This explained that measuring editing efficiency in bulk HSPC cultures underestimates functional graft editing. Across all replicates, engraftment and lineage potency was largely comparable in recipients of edited versus unedited grafts and persistence of CD33 P67 edited cells was on average more than 75%, except in the case of CB 917, around 50% with significant variability (**suppl. Fig. 5** and **suppl. Table 5**).

**Figure 5.**
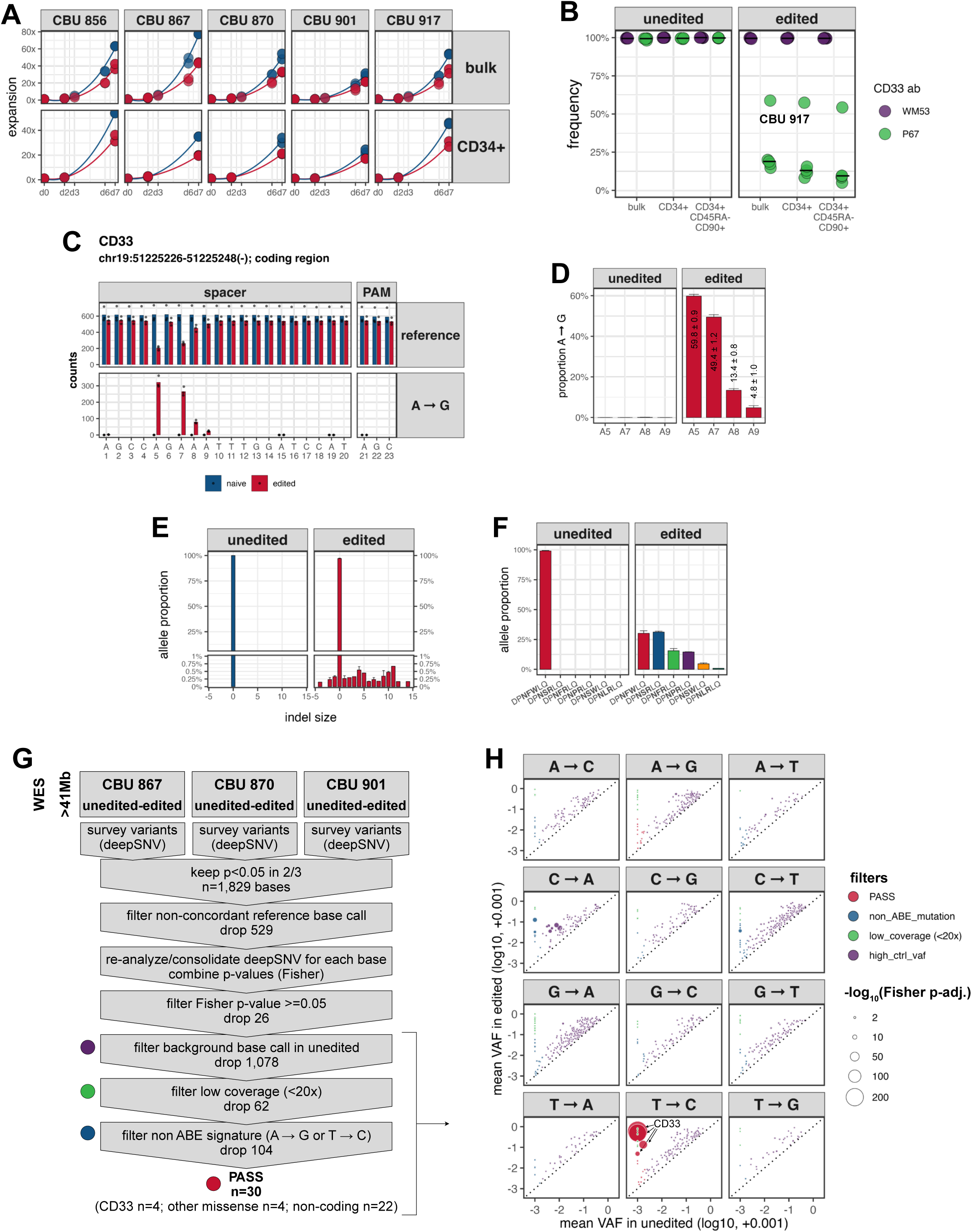
Reproducibility and Safety of CD33 P67 Epitope–Engineered Grafts. A: Expansion kinetics of paired naïve (blue) and CD33 P67 base-edited (red) cord blood (CB) CD34⁺ HSPCs from five donors (CBU 856–917) expanded in UM171-supplemented cultures. Cells were counted on the indicated days, with flow cytometry performed on days 0, 2, and 7. Fold expansion is shown relative to day 0; individual replicates and running averages are plotted. **B:** Day 7 flow cytometry analysis of cells from panel A, stained with CD33 P67 and CD33 WM53 antibodies to assess epitope disruption. Frequencies of antibody-positive cells are shown; CBU 917 exhibited lower editing efficiency. Crossbars indicate median values across five replicates. **C–H:** Whole exome sequencing (WES) analysis of three independent naïve/edited CB units (CBU 867, 870, and 901). **C:** Read counts (PCR duplicates removed, Q >25) within the sg_W22_as-targeted CD33 region. **D:** Frequencies of A-to-G conversions at four adenines within the editing window. **E:** Summary of indel formation across replicates. Error bars represent standard error. **F:** Mean allelic proportions of edited variants derived from WES reads overlapping the CD33 P67 epitope encoding region. Mutant peptides spanning amino acids 18–24 are shown. Additional low-frequency variants are listed in Supplemental Table 7. Error bars denote standard error across replicates. **G:** Exome-wide SNV detection and filtering pipeline. Single nucleotide variants (SNVs) were identified across >41=:Mb of Agilent SureSelect-captured WES regions in three independent pairs of unedited and CD33 base-edited cord blood CD34⁺ HSPCs. The schematic outlines the analysis and sequential filtering strategy. SNV candidates shown in panel **H** (colored dots) represent variants retained prior to the final three filtering steps. Dot color indicates filtering category, and size reflects combined significance across the three donor pairs (–log_10_ adjusted Fisher’s p-value). See Supplemental Table 9 for full candidate list, replicates, counts, and statistical details. VAF, variant allele frequency.

We next explored genomic effects of the CD33 F21/W22 targeting sgRNA in combination with SpG ABE8e (Cas9 R691A) in three donors (CB 867, 870 and 901). To this end, we carried out ultra-deep (200x median coverage) whole exome sequencing (WES) of control expanded (unedited) versus base-edited cultures (**suppl. Table 6**). In these datasets, we observed highly reproducible A-to-G transitions in the editing window of the sgRNA targeted region. Specifically, protospacer positions A5, and A7-9 were edited between 5-60% efficiency, while no significant editing was detected in nearby adenines (**Fig. 5C and D**). Indel formation around the target site was around 3.0±0.6% (**Fig. 5E**, **suppl. Table 7**), in line with original ABE8e reports^34^. Profiling of coding changes based on WES reads spanning the targeted region revealed a ∼70% loss of CD33 WT alleles (red bars in **Fig. 5F**), in exchange for CD33 variant encoding alleles, primarily N_20_S_21_R_22_, N_20_F_21_R_22_, N_20_P_21_R_22_, N_20_S_21_W_22_, and N_20_L_21_R_22_ (**Fig. 5F** and **suppl. Table 7**).

*In silico* off-target predictions^45,46^ identified one candidate locus with two mismatches, 62 loci with three, and 781 loci with four mismatches, of which only 19 localized to coding regions covered by whole-exome sequencing (**suppl. Table 8**). Using deepSNV^47^, a binomial model-based algorithm optimized for sensitive and specific detection of low-frequency variants in deep sequencing data, we interrogated all predicted target and off-target regions also covered by WES (n = 442 bases) and detected no significant A-to-G mutations in any replicate other than those on-target (p-values < 0.0001, **suppl. Fig. 4** and **suppl. Table 8**).

We then extended the deepSNV analysis exome-wide (n=41,621,960 positions including out-of-target regions) to identify cryptic single-nucleotide variants recurrently enriched in base-edited samples (**Fig. 5G**). An initial screen identified 1,829 SNVs that were significant by unadjusted p-value in at least two of three donors. For 529 of these locations deepSNV and database reference base calls were inconsistent and were therefore dropped. After statistical integration of the remaining candidate loci in the three replicates, we applied sequential filtering to exclude non-significant (non-adjusted combined Fisher p-value >=0.05) variants (n=26), positions with background signal in unedited samples (n=1,078), low sequencing coverage (<20, n=62), and variants inconsistent with ABE activity (i.e., not A-to-G or T-to-C, n=104). This resulted in retention of the four on-target and 26 additional candidate positions for downstream validation (**Fig. 5H**, red dots, **5G** and **suppl. Table 9**). Of these, nine mapped to the USP17L gene cluster on chromosome 4p16.1 (**suppl. Fig. 7**), a region of duplicated genes with high sequence similarity known to present read mapping challenges, likely resulting in false variant calls. Among the remaining 17 candidates (**suppl. Fig. 8**), only four were *bona fide* missense variants: *JAK2 N673S* (**suppl. Fig. 6A**) and *POMGNT2 M67T* (**suppl. Fig. 8B**), both annotated in dbSNP (rs1487709003 and rs2089852276, respectively), and two previously undocumented variants, *ZNF83 C515R* (**suppl. Fig. 8C**) and *PRUNE2 S395P* (**suppl. Fig. 8D**).

Since cryptic off-target sites with no obvious protospacer homology are very rare^48^, additional validation and follow-up analyses is warranted in these cases. Of note, none of these four missense off-target candidates were significant by FDR-adjusted combined p-values, nor did they fall into regions with any sgRNA homology.

In conclusion, our data suggest very limited off-target activity in the tested context and support the high fidelity of ABE editing in primary human blood stem cells, particularly when incorporating off-target mitigating mutations such as Cas9 R691A in the base editor design.

## Discussion

Targeted immunotherapies have the potential to transform the treatment landscape for acute myeloid leukemia (AML), yet their clinical utility remains constrained by shared antigen expression on healthy hematopoietic cells. This is exemplified by dose-limiting safety concerns of agents such as gemtuzumab ozogamicin (GO)^49^. Here, we present a clinically scalable base editing strategy that precisely disrupts the CD33 epitope recognized by GO, enabling engineering of immunotherapy-compatible HSPC grafts while preserving CD33 expression and mitigating the risks of a complete gene knockout.

We identified a discrete epitope, surrounding phenylalanine 21 (F21), essential for CD33 P67 antibody binding. Base editor disruption of this epitope abrogated GO binding while preserving CD33 surface expression and ligand-binding capacity, suggesting that epitope editing can decouple therapeutic recognition from endogenous protein function in this case. This approach contrasts with conventional CRISPR nuclease targeting of the CD33 gene^23^, which significantly impairs HSPC expansion and multilineage engraftment due to genotoxicity from double-strand breaks^50^ impacting long-term function of edited cells^51–53^. While CD33’s biological relevance in humans, especially in stem and progenitor cells^54^, remains unclear, we propose that epitope-preserving editing may offer a more physiologic alternative to gene ablation, balancing therapeutic protection with functional conservation.

Importantly, our strategy was integrated into an optimized *ex vivo* manufacturing platform using UM171-mediated expansion of cord blood-derived CD34⁺ HSPCs. UM171 supports the expansion and rejuvenation of immature, repopulating HSPCs during culture, significantly improving the clinical applicability of cord blood grafts. In this context, epitope editing can be achieved at high efficiency, and retains robust *in vivo* graft performance, with balanced lympho-myeloid output and durable T-cell potential. Notably, engraftment levels were maintained even under high dose GO challenge, confirming effective immunologic shielding of the graft. This highlights the unique synergy between precise epitope editing and UM171-based expansion in enabling safe and functionally intact HSPC graft engineering.

From a translational perspective, this approach offers several advantages. First, the use of base editors avoids the cytotoxicity associated with nuclease-mediated editing. Second, the precision of the epitope edit allows selective immune evasion while preserving potentially beneficial CD33 functions. Third, since CD33 is expressed widely in the healthy hematopoietic system, editing the CD33 P67 epitope will drastically reduce GO target sites competing with residual malignant cells in transplanted patients. We hypothesize that this will lower the minimal effective dose for AML cell eradication, especially during early phases after MRD detection post-transplant. Fourth, the reproducibility of editing across multiple donors, the observation that editing is high in immature subsets and that incomplete editing is still effective in protecting against myelosuppression, supports feasibility for a clinical manufacturing strategy. While we cannot fully exclude subtle effects on CD33’s immune regulatory roles, our data suggest that key signaling and internalization functions are retained post-editing, and ongoing work will clarify these effects in inflammatory or disease-relevant contexts.

More broadly, the ability to engineer therapeutic resistance into HSPC grafts during UM171-supported *ex vivo* culture without compromising engraftment is critical for integrating targeted immunotherapies into the post-transplant context. As editing technologies and specific immunotarget editing strategies will evolve further, the convergence of gene editing, *ex vivo* stem cell expansion, and immunotherapy promises to reshape how we approach relapse prevention in hematologic malignancies.

## Methods

### CD34^+^ cord blood cell culture

Human CB-derived CD34⁺ cells were cultured in hematopoietic stem and progenitor cell (HSPC) expansion medium containing SCGM (CellGenix, #20806-0500), supplemented with 100 ng/mL SCF, 100 ng/mL Flt3L, 50 ng/mL TPO, and 50 ng/mL IL-6 (Shenandoah Biotechnology), 35 nM UM171 (ExCellThera), and Gentamicin (Gibco, #15750-060). Cell concentrations (viable total nucleated cells, vTNC/mL) were monitored using Via1-Cassettes (ChemoMetec, #941-0012) during culture.

### mRNA production

Synthesized DNA fragments encoding 5′ and 3′ untranslated regions (UTRs) and coding sequences (IDT; Twist Bioscience) were cloned into a pUC19 backbone via Gibson assembly. For in vitro transcription (IVT), linearized plasmids (EcoRI) were PCR-amplified using a forward primer introducing a CleanCap-AG-compatible T7 promoter and a reverse primer encoding a 120-nucleotide poly(A) tail. PCR (NEB Q5, M0491) was performed over 30 cycles using 50 ng template in four 50-µL reactions. Amplicons were purified (GeneJET, K0702), yielding ∼400 ng/µL in 40 µL nuclease-free H_2_O. IVT was performed using 1 µg of PCR product and the HiScribe T7 mRNA Kit with CleanCap AG (NEB, E2080S), substituting UTP with N1-methyl-pseudouridine-5′-triphosphate (Jena Bioscience, NU-890L). mRNA was column-purified (NEB, T2050), quantified by Nanodrop (typical yield ∼2 µg/µL), and assessed for integrity (Bioanalyzer, IRIC). Final products were aliquoted and stored at −80°C.

### CD33 sgRNA library screening

A sub pool of a 119/120-nt oligonucleotide library (Twist Bioscience) containing 128 unique sgRNAs targeting CD33 exon 2 was PCR-amplified to pre-saturation using Q5 High-Fidelity 2X Mastermix (NEB, #M0492L). Amplicons included primer-encoded homology arms for Gibson Assembly (GA) into an Esp3I-digested, dephosphorylated pLKO5-based lentiviral vector containing an EFS-tRFP657-P2A-Puro cassette (derived from Addgene #57824). PCR products were purified, quantified, and quality checked. GA was performed using 2x NEBuilder HiFi mix (NEB, #E2621L) at a 2:1 insert:vector molar ratio. Reactions were transformed into NEB Stable Competent E. coli (NEB, #C3040I). 1% was plated for colony counts; the remainder was expanded in liquid culture, and plasmids were purified by Midiprep (PureLink HiPure, ThermoFisher, #K210005). Estimated library coverage was ∼780x. Cloned libraries were validated by restriction digest and next-generation sequencing (Illumina). OCI-AML5 cells were transduced at MOI <0.2 to ensure single-copy integration, confirmed by tRFP657 FACS analysis. Cells were selected with 1 µg/mL puromycin prior to nucleofection (1*10^6^cells) with 6 µg of SpRY-ABE8e or SpRY-TadCBEd mRNA. sgRNA representation was assessed in plasmid and genomic DNA before and after nucleofection, and in sorted de-epitoped populations to identify target sequences. For this, the guide sequence was amplified with Q5 High-Fidelity 2x Mastermix (NEB # M0492L) and primers with TruSeq adapters and indices. NGS amplicon-seq was performed on an Illumina NextSeq 2000 (IRIC Genomics Platform) with a minimum library coverage of 1000x. sgRNA enrichment was calculated based on library size-normalized counts (MAGeCK^55^) following bowtie^56^ read alignment. Editing-induced mutations in sorted populations were identified by PCR-amplified CD33 exon 2 followed by Nanopore amplicon sequencing (Plasmidsaurus).

### CD34^+^ cell editing

CD34⁺ cells were thawed and cultured for 48 h in HSPC expansion medium (2.5*10^5^ cells/mL) containing UM171 (35 nM). Nucleofection was performed using the Amaxa 4D-Nucleofector Core and X units (Lonza, #AAF-1002B, #AAF-1002X) with P3 Primary Cell Buffer (Lonza, #CA10064-500) and 100 µL cuvettes (Lonza, #V4XP-3024) using program CA137. Cells (1*10^6^ per 100 µL reaction) were nucleofected with 5 µg each of Cas9 or base editor mRNA and sgRNA (Synthego) for the 1x condition, or 10 µg each for the 2x condition. Post-nucleofection, cells were cultured at 2.5*10^5^ cells/mL. Media volume was adjusted daily to re-establish post-nucleofection density. Transplantation was performed on day 5 post-nucleofection (day 7 of culture).

### Mylotarg treatment

4.5 mg GO/Mylotarg (Pfizer) was aliquoted as lyophilized powder and stored at 8°C. The active compound-to-carrier ratio was determined to be 1:32. On the day of use, aliquots were freshly reconstituted by addition of sterile H2O to reach a concentration of 1 μg/ul of active GO. This solution was then diluted tenfold in PBS and kept on ice, protected from light until injected in mice. Administration was done by tail vein injection and dosed at 1.0 mg/kg. PBS was injected to control cohorts.

### Human CD34+ cord blood cells

Cord blood collection and use were approved by the Research Ethics Boards of the University of Montreal, Maisonneuve-Rosemont Hospital, and Charles-LeMoyne Hospital (QC, Canada). Umbilical cord blood units were obtained with informed consent at Charles-LeMoyne Hospital.

CD34⁺ cells were isolated using the EasySep Human Cord Blood CD34 Positive Selection Kit II (StemCell Technologies, #17896). Cord blood was incubated with RosetteSep cocktail (5 µL/mL, 20 min, RT), diluted 1:1 with separation buffer (PBS, 2 mM EDTA, 2% FBS, 1 µL/mL DNase), and layered (max. 30 mL) onto 15 mL Lymphoprep (StemCell Technologies, #07851) for density gradient centrifugation (20 min, 1200 × g, no brake). Mononuclear cells were harvested, washed (300g, 10 min, low brake), and resuspended in separation buffer. CD34⁺ selection was performed per manufacturer’s protocol. Cells were used immediately or cryopreserved in FBS with 10% DMSO.

### Cell line culture

Culture of HEK293T, AML5 and U937 cells was done in DMEM (Gibco, Cat#12491-015), aMEM (Gibco, Cat#12751-063), RPMI 1640 (Gibco, Cat#22400-089), respectively, supplemented with 10% Heat-inactivated Fetal Bovine Serum (Wisent, Cat#098-150) and 10ng/mL GM-CSF (Shenandoah, Cat#100-72) for AML5 cells.

### Cell Line Editing

Cell line editing was performed using the same conditions as for CD34⁺ cells, except for 1M electroporation buffer (5 mM KCl, 15 mM MgCl₂, 120 mM Na₂HPO₄/NaH₂PO₄, pH 7.2, 50 mM mannitol). For nucleofection in 2×8 20 µL cuvette strips (Lonza, #V4XP-3032), 2.0*10^5^ cells were electroporated with 1 µg mRNA and 1 µg sgRNA. Conditions were scaled proportionally for 100 µL cuvettes. Post-nucleofection, cells were transferred to culture medium containing gentamicin, expanded, and FACS analyzed or sorted.

### Flow Cytometry

Fresh or cultured cells were analyzed using a BD FACSCanto II (panels 11 and ISHAGE_CD3) or BD LSRFortessa (other panels) and sorted using a BD FACSAria II (BD Biosciences). Cells were processed in FACS buffer (PBS, 2% FBS, 2 mM EDTA, ±100 µg/mL DNase). For HSPC phenotyping, cells were stained using antibody panels 7, 8, 11, and ISHAGE_CD3 (Supplemental Table S3). FACS data were analyzed using FlowJo v10 and plotted in R.

### Transplantation and Engraftment Analysis

Day 7 expanded CB-derived CD34⁺ HSPCs were transplanted via tail vein into sublethally irradiated (250 cGy) 8–16-week-old female NSG mice (NOD-Scid IL-2R=: null; The Jackson Laboratory). Cell doses and engraftment data are detailed in the corresponding figures and supplemental tables. Engraftment was assessed by flow cytometry at defined timepoints using bone marrow (BM) cells collected via femoral aspiration (early timepoints) or flushing of femurs, iliac bones, and tibiae (final timepoint). Spleen and thymus were also analyzed at endpoint. Samples were processed using in-house RBC lysis buffer, washed, blocked, and stained with antibodies listed in the supplemental methods table. All animal procedures complied with recommendations of the Canadian Council on Animal Care and were approved by the Deontology Committee on Animal Experimentation at the University of Montreal.

### Synthesis of CD33L-PEG-DSPE

Chemical synthesis is described in detail elsewhere^42^.

### Liposome Preparation

Naked and CD33L liposomes were prepared as described^42^. Stock solutions of DSPC (10 mg/mL), cholesterol (5 mg/mL), DSPE-PEG (4 mg/mL), AF647-PEG-DSPE (1 mg/mL), and CD33L-PEG-DSPE (3.2 mg/mL) were used. Lipid components were mixed in chloroform to achieve final molar ratios of 57:38:4.8:0.1 (DSPC:cholesterol:DSPE-PEG:AF647-PEG-DSPE 7) for naked liposomes, and 1 mol% CD33L-PEG-DSPE was added for CD33L liposomes. Solvent was evaporated under nitrogen gas, and 100 µL DMSO was added to resuspend the dried lipids. CD33L-PEG-DSPE and AF647-PEG-DSPE were added from frozen DMSO stocks (−80°C). Samples were cryo-lyophilized overnight, and stored at-80°C. For extrusion, dried lipids were rehydrated in PBS (pH 7.4), sonicated in 1-min on/5-min off cycles until uniformly suspended, and extruded sequentially through 800 nm and 100 nm filters. Final liposome size (∼120 ± 20 nm) was confirmed by dynamic light scattering (Zetasizer Nano S, Malvern Panalytical). Liposomes were shipped and stored at 4°C.

### CD33 ligand binding studies

All U937 cell conditions were maintained at equal densities one day before the assay: 0.2*10^6^ cells/mL in a T25 flask. Cells were collected and resuspended in 50uL fresh media in 96-well U-bottom plate at 100,000 cells/well. 50uL of media containing naked or CD33L conjugated liposomes (labeled with AF647, at 1 mol% 1mM, diluted at 1/20) was added per well. Cells and naked / CD33L liposome suspensions were incubated for 1h at 37C. Following incubation, cells were washed once before blocking (10min) and CD33 HIM3-4 staining (20min) at room temperature. Two final washes in FACS tubes were performed prior to flow cytometry acquisition.

### Whole Exome Sequencing (WES)

CD34+ cells from three separate cord blood donors were cultured in UM171 for 7 days ± d2 ABE8e/sgW22_as editing as described above. 5*10e6 cells were collected, washed and genomic DNA was extracted (QIAGEN, DNeasy Blood & Tissue kit, 69504). Exome sequencing was done using the SureSelect Human All Exon V8 panel (Agilent, 5191-6873) followed by paired-end sequencing (Illumina NovaSeq 2500M, read length 2x 150) to equal depth (1000x) in all six samples. Sequences were trimmed for sequencing adapters and low quality 3’ bases using Trimmomatic version 0.38 and aligned to the reference human genome version GRCh38 (gene annotation from Gencode version 32, based on Ensembl 98) using BWA version 0.7.17.

Samtools version 1.19 was then used to fix mate coordinates and Picard used to mark read duplicates.

### Variant Detection and Statistical Analysis

To identify single-nucleotide variants (SNVs) introduced by base editing, we analyzed three biological replicate pairs of control and base-edited cord blood CD34⁺ cells using the deepSNV R package^47^. This method is specifically designed to detect low-frequency variants in paired deep-sequencing data by modeling error rates from a control sample. Non-ambiguous, PCR-duplicate-excluded and high base call quality filtered allele counts were first extracted using the bam2R function (mask = 3328, q = 25) across 41,621,960 genomic positions. A binomial model was employed, with the unedited sample serving as the control to estimate site-specific error rates and to call reference alleles. Statistical testing was performed with the alternative hypothesis set to “greater”, detecting increases in variant allele frequency in the edited sample relative to the control. Each replicate was first analyzed independently in 114 data chunks to nominate candidate loci with significant SNVs. For downstream analysis, we then retained SNVs with p-values < 0.05 in at least two of the three replicate pairs (n = 1,829), ensuring statistical robustness and biological reproducibility. To recover loci with missing p-values (≥ 0.05), deepSNV was re-run and variant summaries, including per-base p-values, allele frequencies, and coverage, were extracted using a custom summary function built around deepSNV output. Results across all replicates were merged, with replicate identifiers embedded for downstream tracking. To integrate statistical evidence across replicates, Fisher’s method (via the metap R package) was used to combine p-values per locus and variant base call. These were further adjusted for multiple testing using the Benjamini-Hochberg (BH) procedure. Missing p-values resulting from no counts were imputed as 1. For each locus, the most significant variant base call by Fisher p-value was retained. Next, we compared the reference base reported by deepSNV to the true genomic base retrieved using the getSeq() function from the BSgenome.Hsapiens.UCSC.hg38 package and filtered loci with discordant reference base calls (n=529). We then excluded loci with (non-adjusted) Fisher p-values ≥ 0.05 (n=26), loci with mean control sample variant allele frequencies >0.1% (n=1,078), and low sequencing coverage (<20x) in either sample (n=62). After applying all filters, 104 loci remained. Of these, we focused on SNVs with mutation signatures characteristic of adenine base editors (ABE), specifically A→G (n = 15) and T→C (n = 15) substitutions, for manual inspection.

## Acknowledgments

The authors thank Melanie Frechette, Valérie Blouin-Chagnon and Koryne Léveillé for assistance with mouse experiments, Annie Gosselin and Angelique Bellemare for assistance in fluorescence-activated cell sorting, and Muriel Draoui for project administration. This project was funded by the Canadian Stem Cell Network through a Horizon Awards Program 2022-2025 entitled ‘Engineered hematopoietic stem cells (eHSCs) as vehicles for next generation therapies’ awarded to G.S., E.G. and others.

## Author Contributions

B.L. conceived the project, designed and performed CD33 epitope mapping experiments, generated mRNA reagents and sgRNAs, conducted liposome binding studies, performed bioinformatic analyses and data interpretation, generated all figures, and wrote the manuscript.

M.L. optimized and carried out CD34⁺ cell editing and transplantation experiments, including phenotypic, expansion, and engraftment analyses, and performed data analysis, conducted liposome binding studies. S.C. assisted in validating the CD34⁺ editing and transplantation pipeline, including phenotypic, expansion, and engraftment analyses, and contributed to data analysis. T.M. assisted with CD33 sgRNA library-based epitope identification. M.J. prepared the CD33 ligand-lipid conjugates. E.N.S. prepared and quality-checked both naked and CD33 ligand-decorated liposomes. J.F.S. assisted with sgRNA library data analysis. N.M., S.M., M.F.T., S.R.S., M.T., C.J.C., and V.D. assisted with NSG tissue preparation. I.B. and L.P. prepared and banked cord blood-derived CD34⁺ cells. J.C., E.T., and E.G. provided essential technical and conceptual expertise. M.S.M. contributed expertise on CD33 biology and the design of functional studies.

G.S. oversaw overall study design, data interpretation, and project supervision.

## Data availability

Whole exome sequencing (WES) data have been deposited in the NCBI Sequence Read Archive (SRA).

---

**Supplemental Figure 1:**
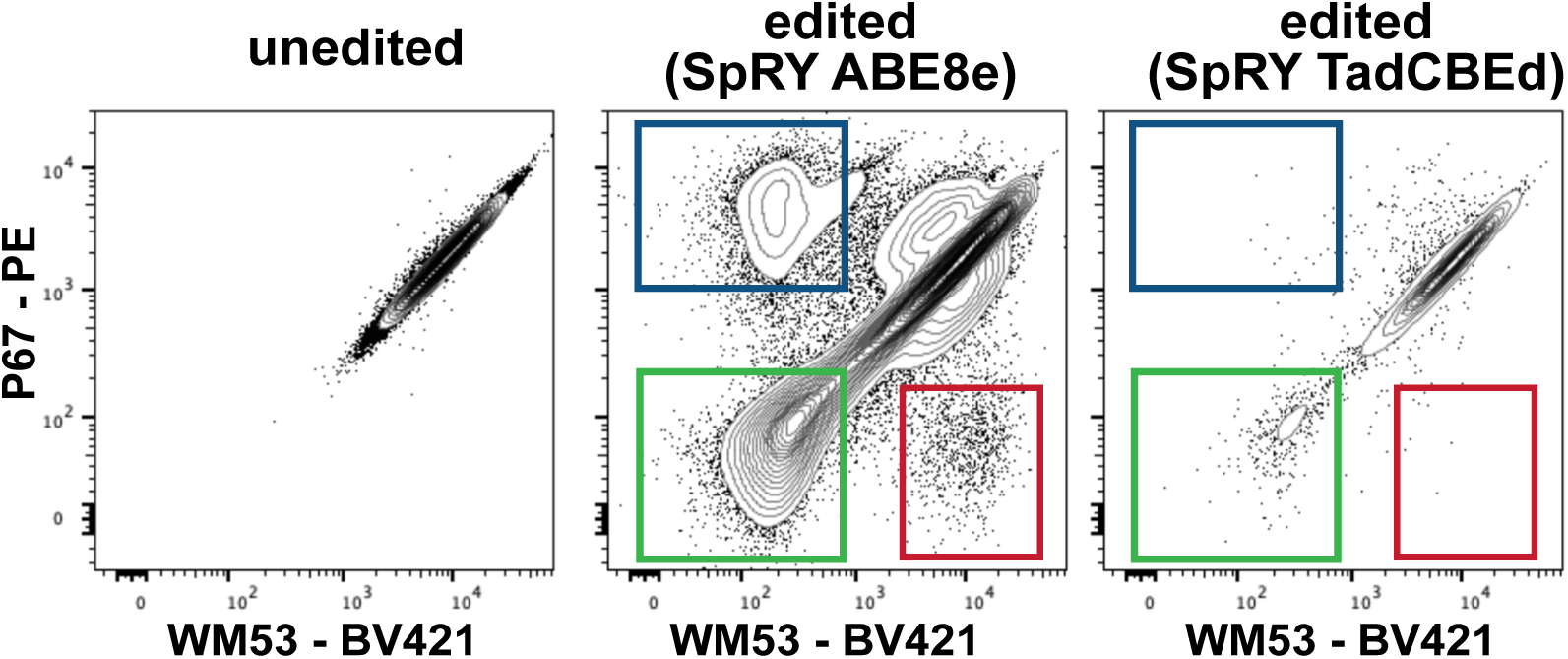
Outcome of TadCBEd in CD33 sgRNA library expressing cells. In comparison to SpRY-ABE8e editing (left and middle is a replicate experiment of Figure 2B), SpRY-TadCBEd electroporation yielded few CD33 P67 or WM53 de-epitoped cells and was thus not further pursued. One of two independent experiments is shown.

**Supplemental Figure 2:**
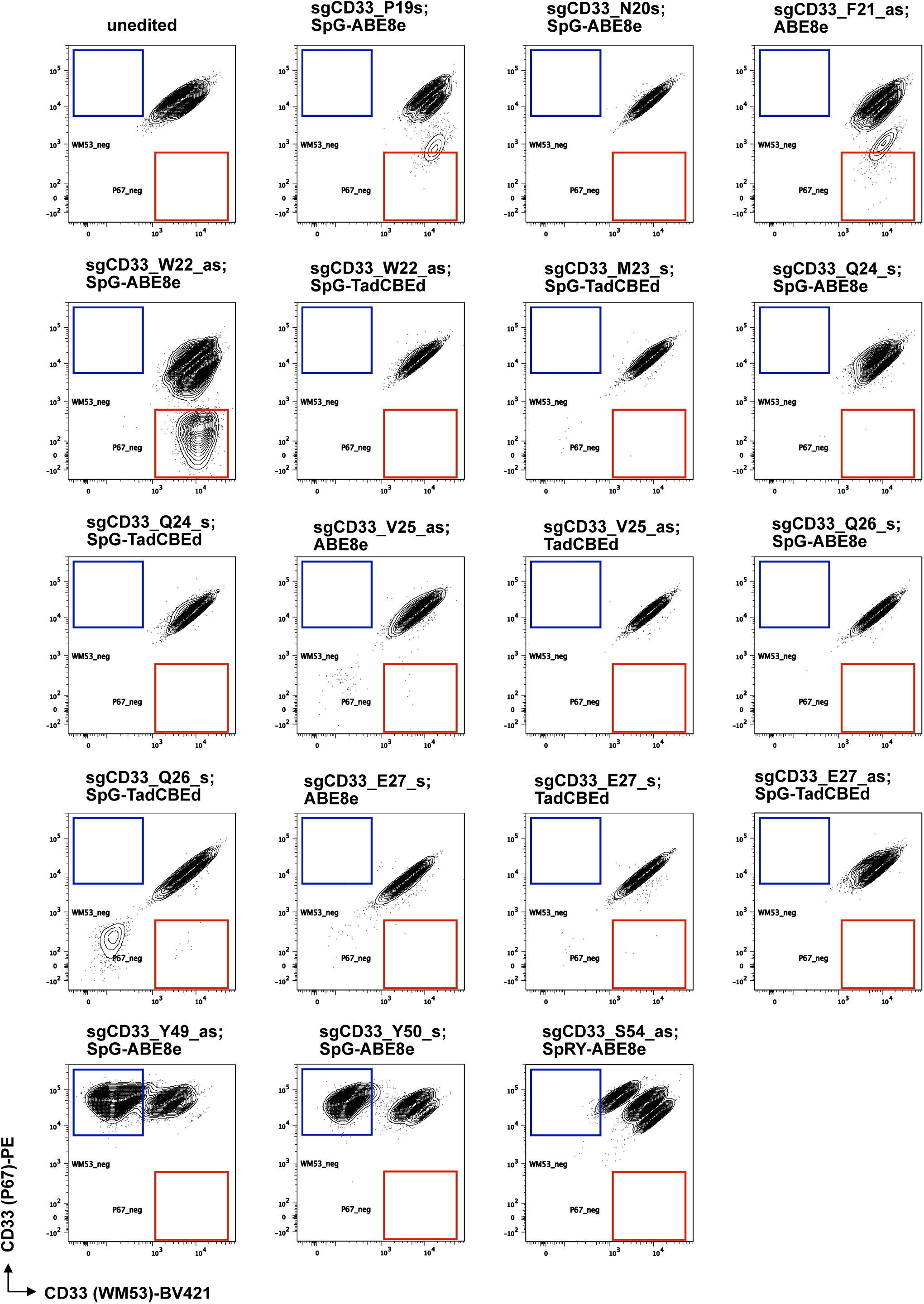
Validation experiments of individual sgRNAs flanking the CD33 P67 and WM53 epitope regions. AML5 cells were electroporated with the indicated combinations of synthetic sgRNAs (Synthego, see **supplemental table 2** for sequences) and PAM-recognition modified ABE8e or TadCBEd encoding mRNAs one week before double staining with CD33 P67 and WM53 antibodies and FACS analysis. Blue and red gates identify CD33 P67 and WM53 de-epitoped cells, respectively. Based on these validation experiments, we selected the sgCD33_W22_as / SpG-ABE8e and sgCD33_Y49_as / SpG-ABE8e combinations as most efficient for CD33 P67 and CD33 WM53 epitope engineering.

**Supplemental Figure 3:**
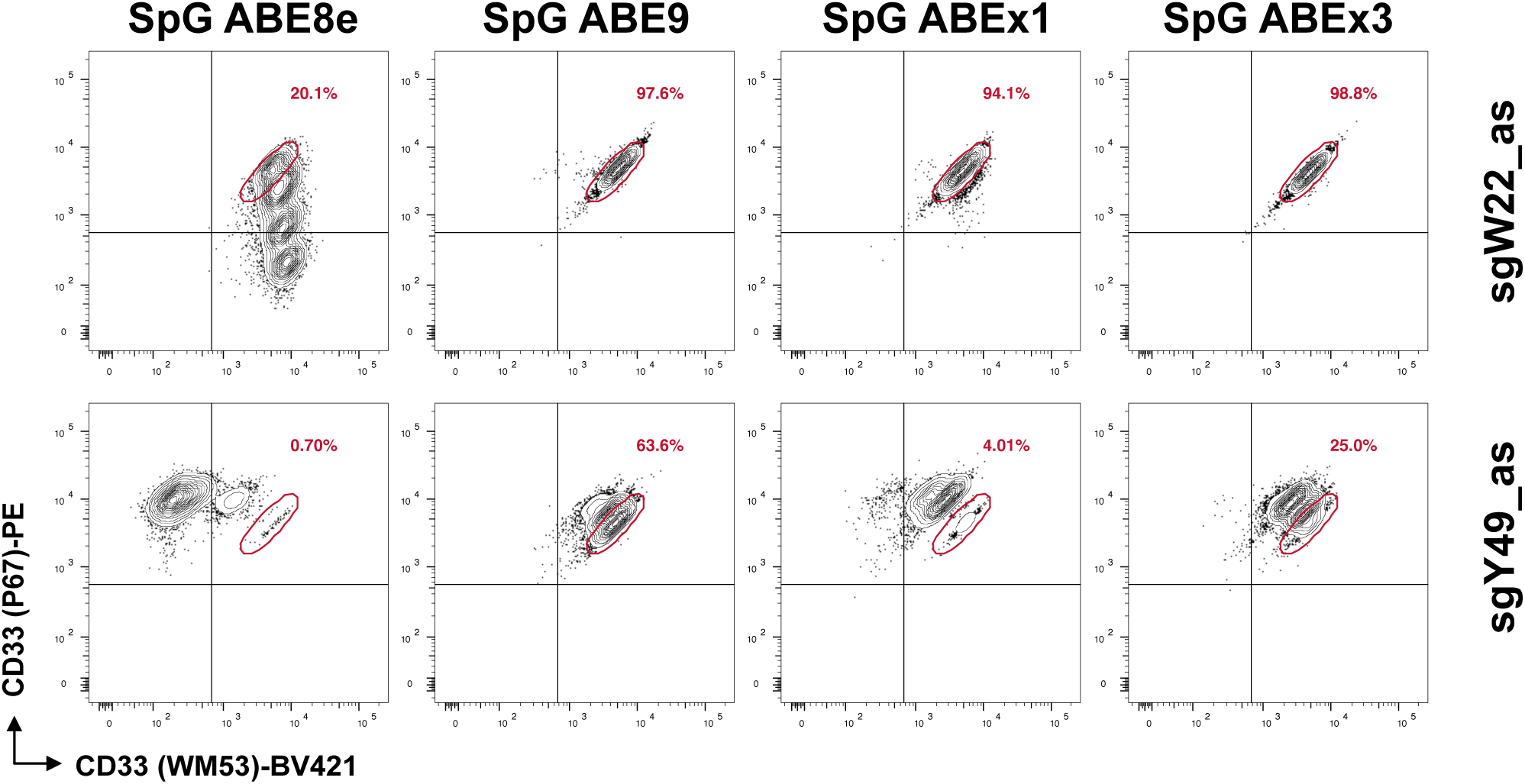
Validation experiments to determine best ABE base editor for CD33 de-epitoping. AML5 cells were electroporated with either sgW22_as or sgY49_as synthetic sgRNAs (Synthego) combined with different SpG (NGN PAM-compatible) ABE variant encoding mRNAs. In comparison to ABE8e, ABEs with narrow editing windows (ABE9, ABEx1 and ABEx3) failed to efficiently generate epitope-disrupted alleles. Cells were FACS analyzed 5 days post-electroporation.

**Supplementary Figure 4:**
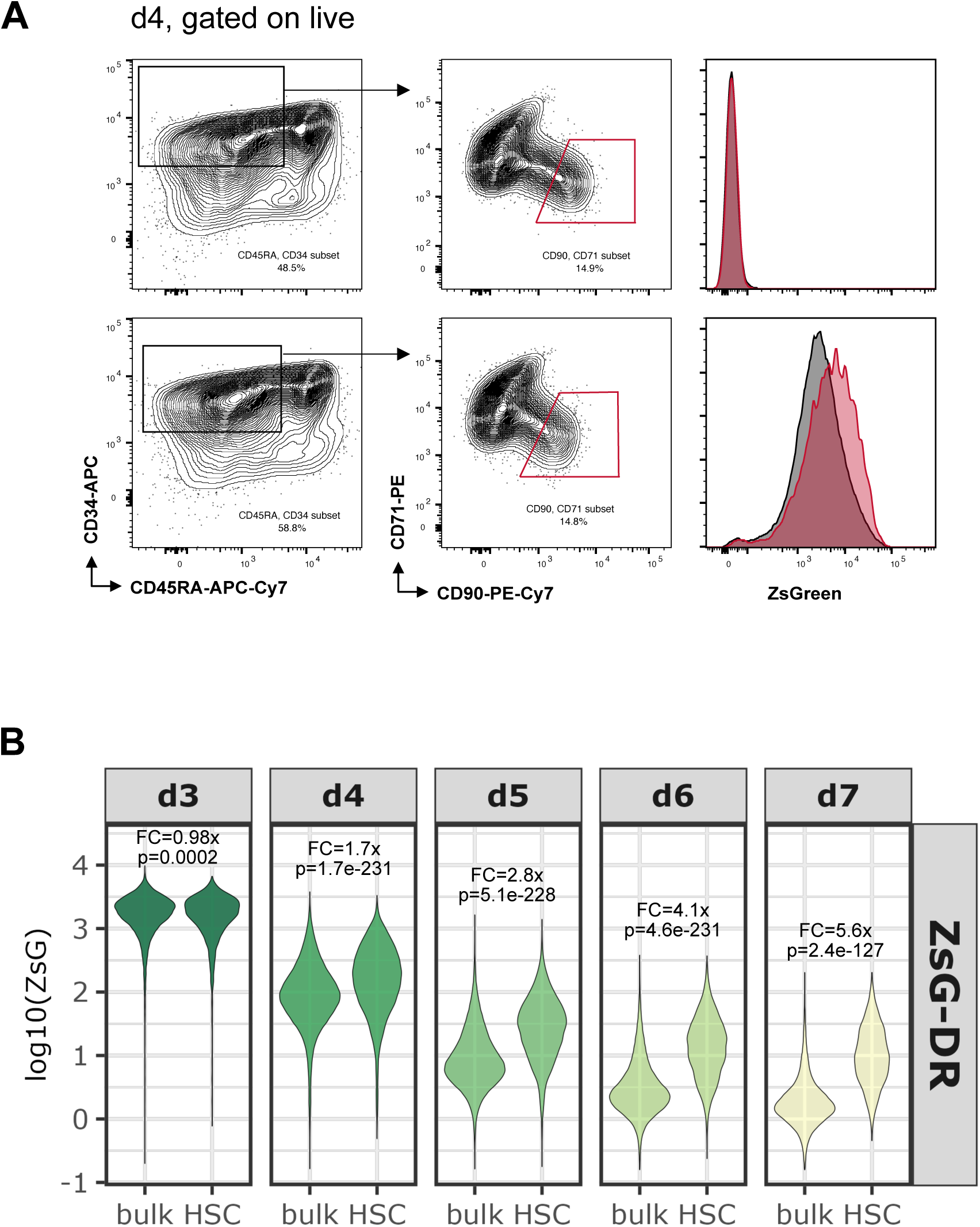
Prolonged mRNA Retention in HSC-Enriched Populations Within CD34+ HSPC Expansion Cultures. After 2 days of culture, cord blood-derived CD34+ were electroporated with mRNA encoding ZsGreen coupled to a human Ornithine De-Carboxylase (hODC) degron (ZsG-DR) (PMID 10611396) for rapid protein degradation. Starting 1 day later (d3), daily FACS analysis using stem cell surface markers CD34, CD45RA, CD90, and CD71 (PMID 38207291) was conducted to assess ZsG signal stability. **A:** Gating strategy employed to measure ZsG signal in bulk versus HSC-enriched subpopulations (CD34+/CD45RAlow/CD90high/CD71low; red gate). **B:** Scaled ZsG fluorescence values (exported from FlowJo) are plotted over time and were compared using a Benjamini-Hochberg corrected two-sided t-test. ZsG-DR signal, which serves as a proxy for its mRNA, exhibits prolonged stability within slower-cycling HSPC populations compared to bulk.

**Supplementary Figure 5:**
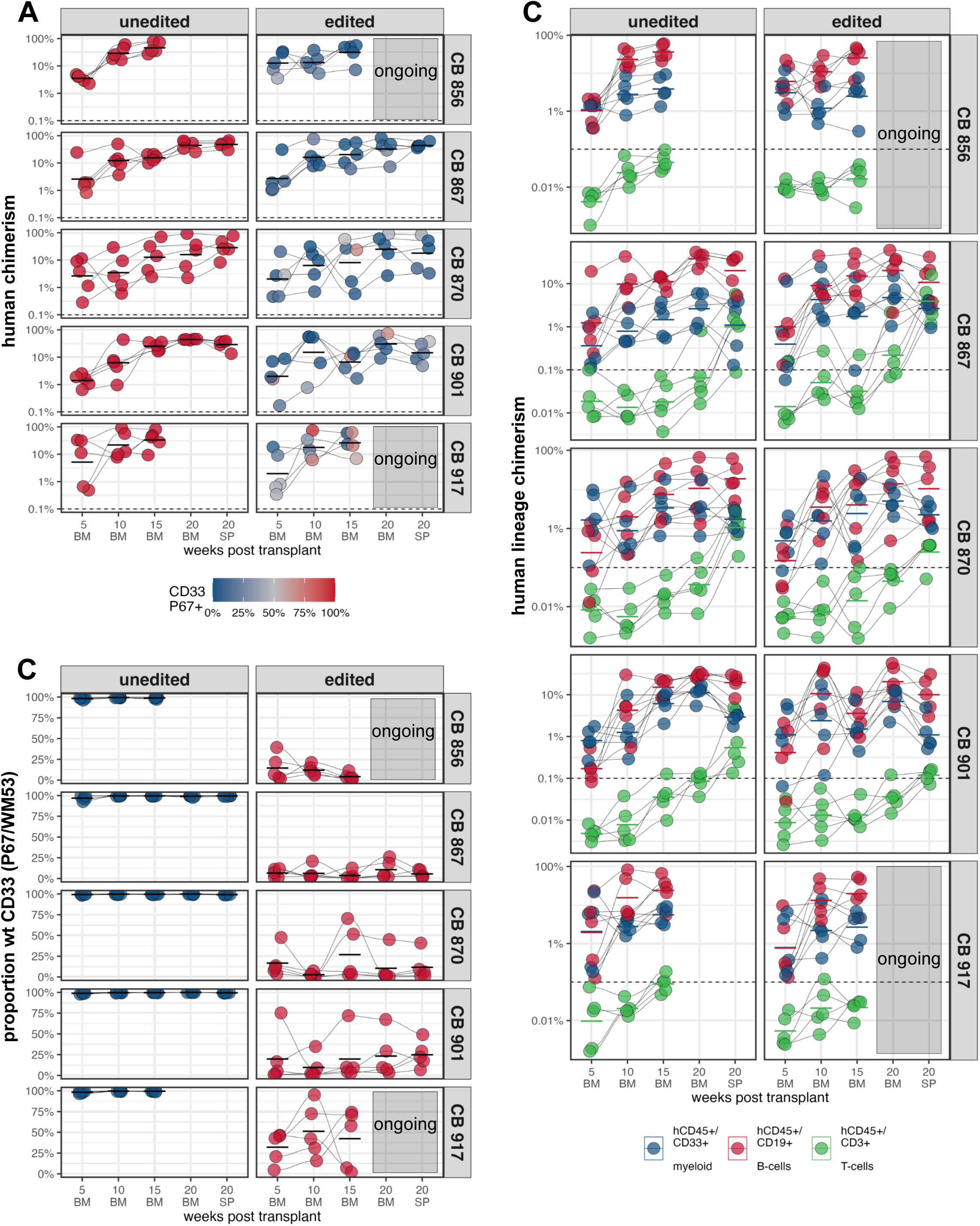
CD33 graft engineering reproducibility. Cells from Figure 5A were transplanted into NSG mice (progeny of 5K CD34+ cells plated at d0). Summary of five unedited versus edited replicate pairs. **A:** Total human engraftment with CD33 P67/WM53 proportions expressed as color-scale. **B:** P67 cross-reactivity on graft-derived CD33^+^ cells plotted as percentage; **C:** human lineage engraftment, separated by color code. Crossbars indicate mean, connecting lines indicate different mice across timepoints. n= 5 mice for each cohort. CB 856 and CB 917 transplantation experiments are ongoing at the time of submission.

**Supplementary Figure 6:**
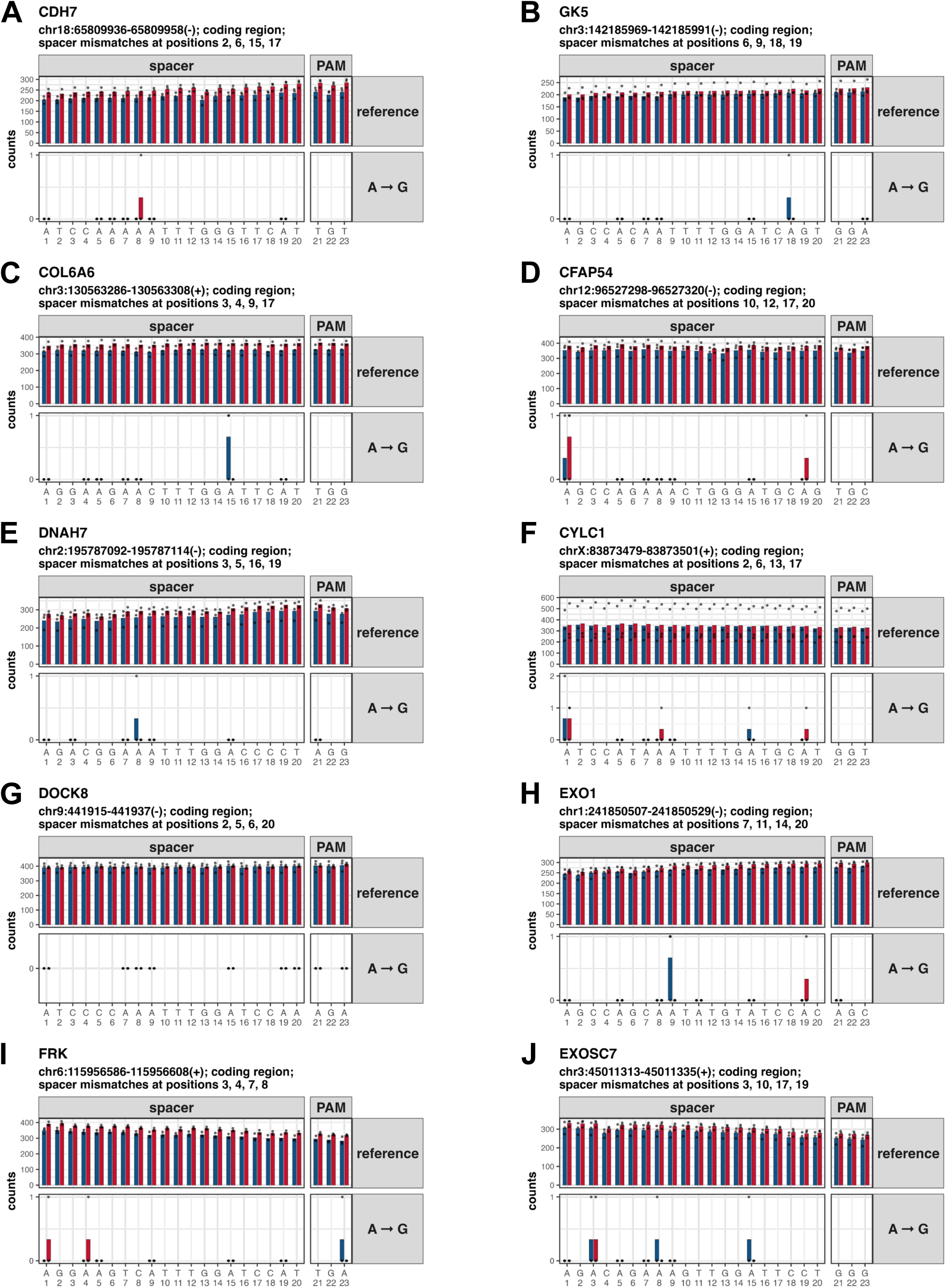
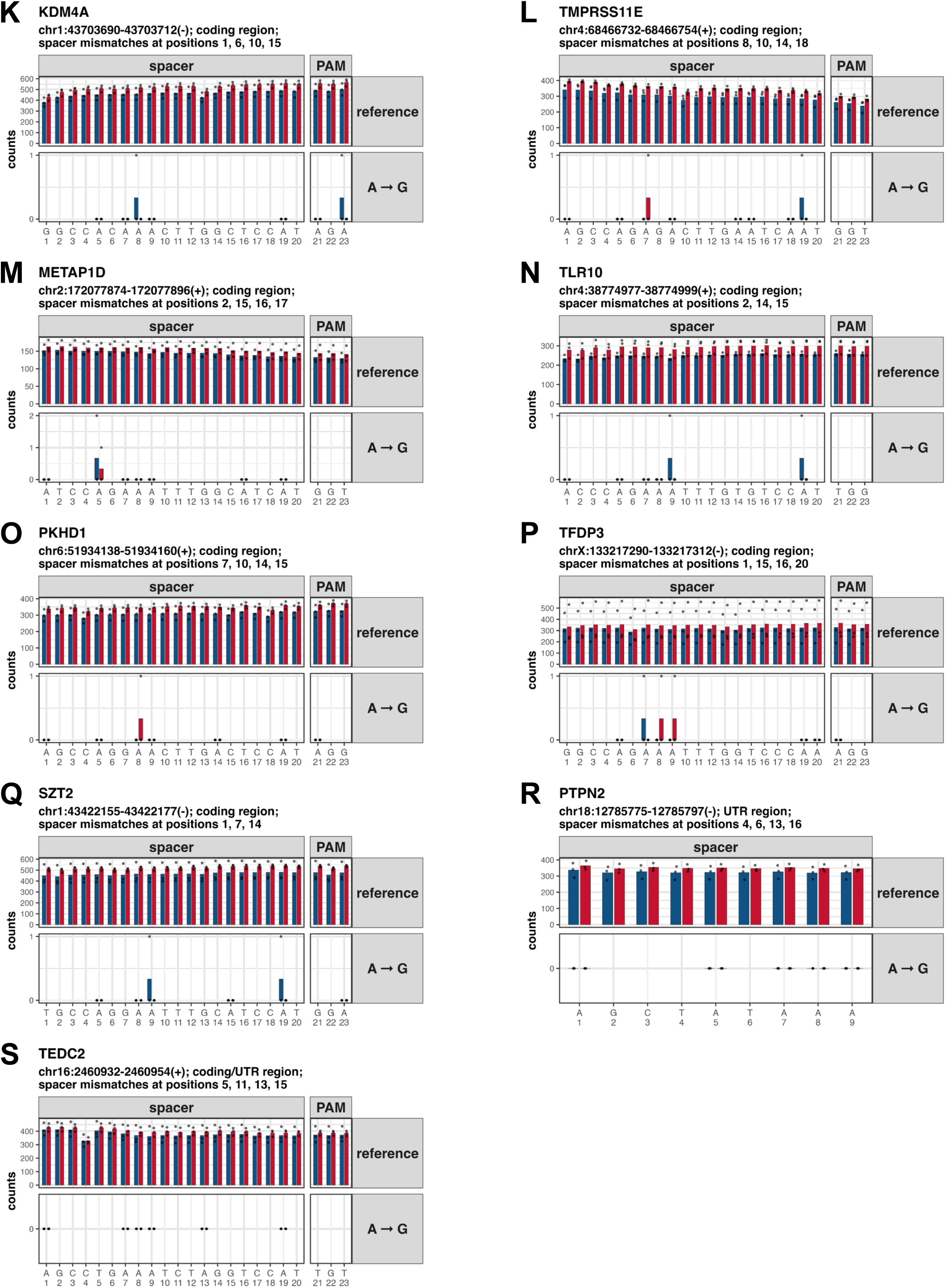
Mutational summaries of WES covered predicted off-targets. Reference base counts (PCR-duplicate excluded and quality-filtered, q>25) are plotted for unedited (blue) versus edited (red) samples across protospacer and PAM regions of predicted off-target regions. For each Adenine, potential Guanine variant base counts are plotted in the A➔G facet. Genomic positions, strand, mismatch positions and overlapping genes are indicated for each position. No significant A➔G mutations were identified in these regions.

**Supplementary Figure 7-8:**
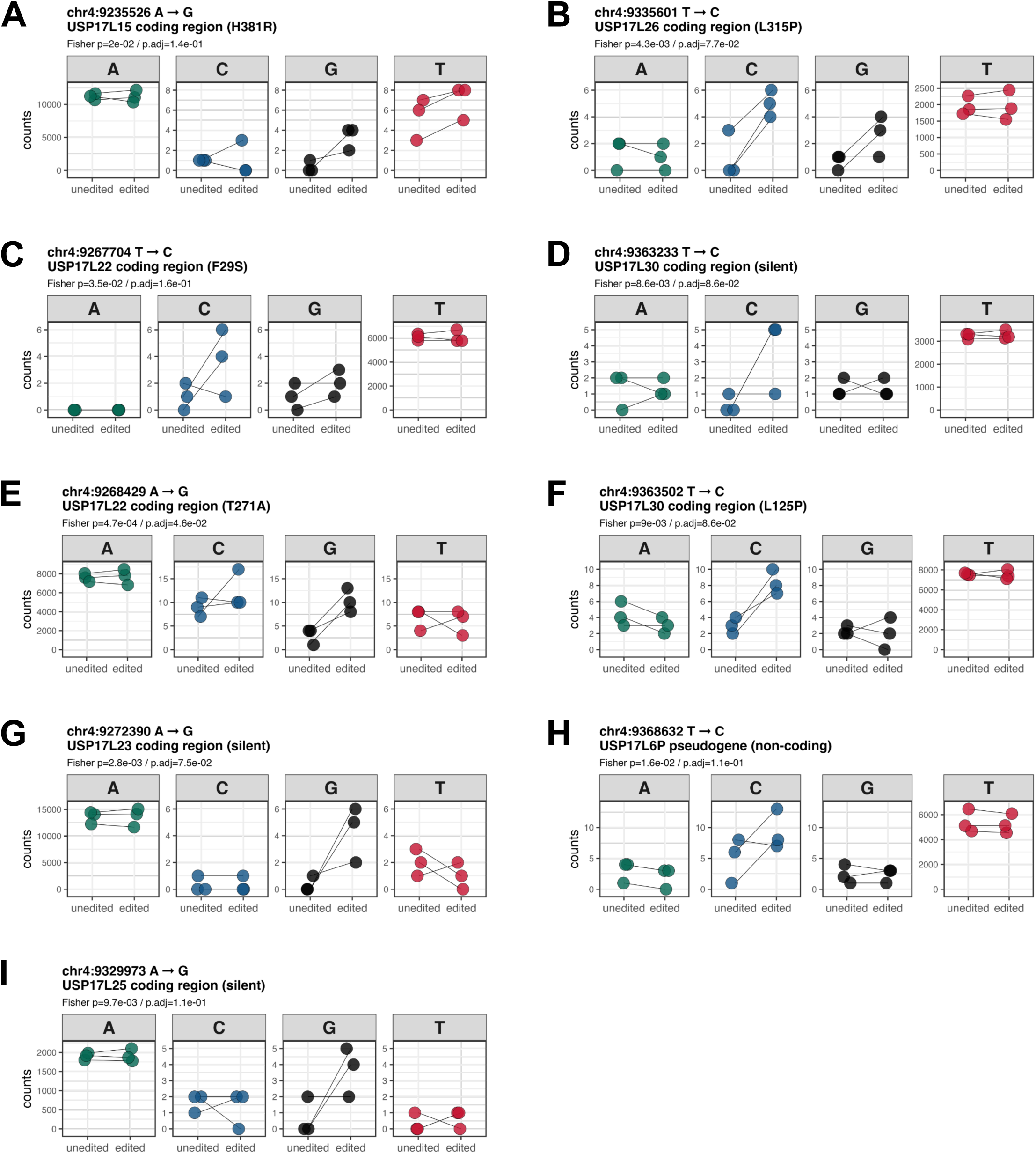
Mutational summaries of SNV candidate positions. Base counts (PCR-duplicate excluded and quality-filtered, q>25) are plotted for unedited (blue) versus edited (red) samples at positions identified in the SNV discovery pipeline outlined in Figure 5G. Genomic positions, ABE compatible mutation patterns (A➔G or T➔C), overlapping features and combined p-values are indicated for each position. Positions in **Supplementary Figure 7** locate to the USP17L gene duplication cluster on chromosome 4; positions in **Supplementary Figure 8A-D** result in missense mutations, whereas positions in **E-P** are silent or non-coding mutations.

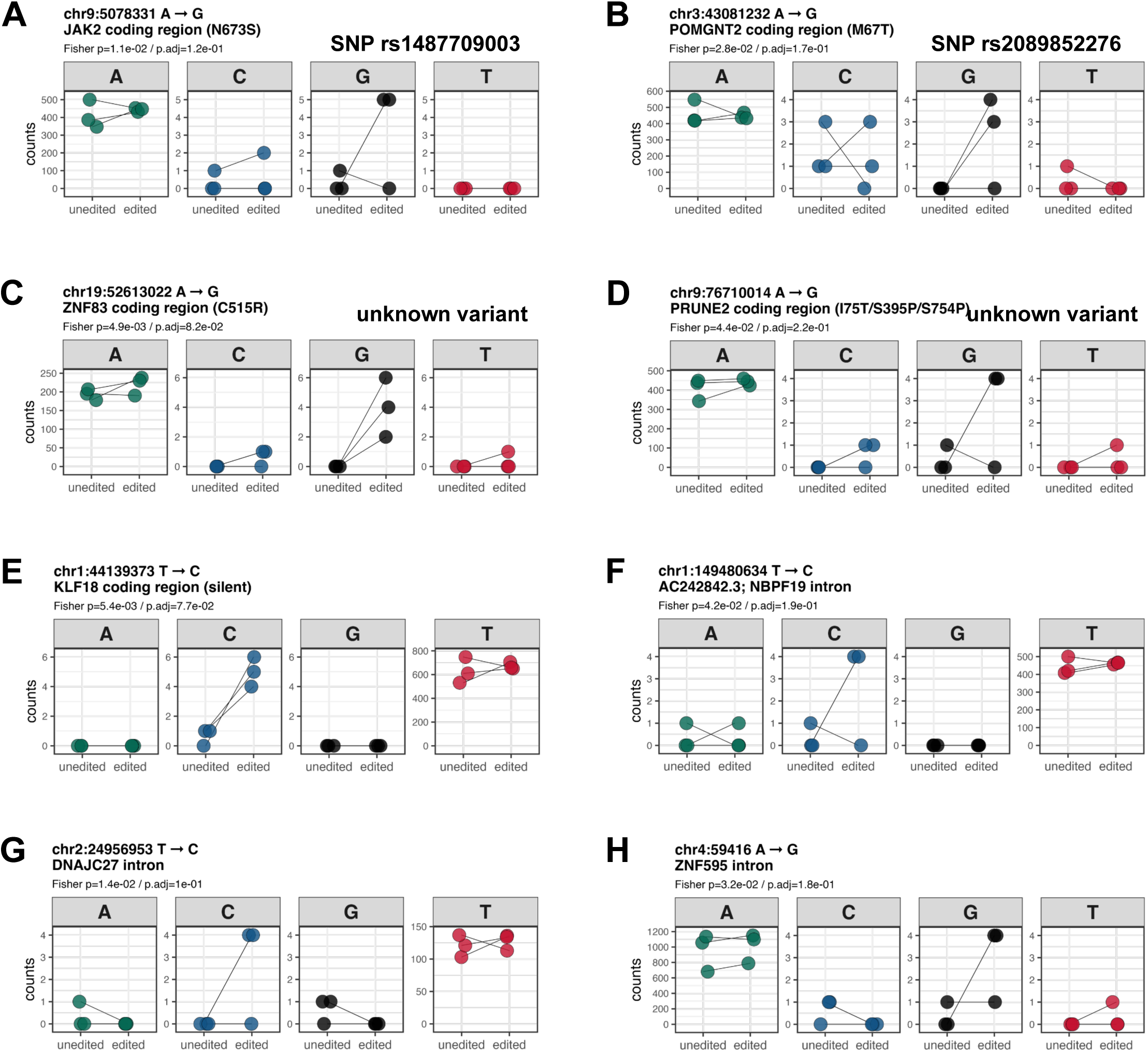
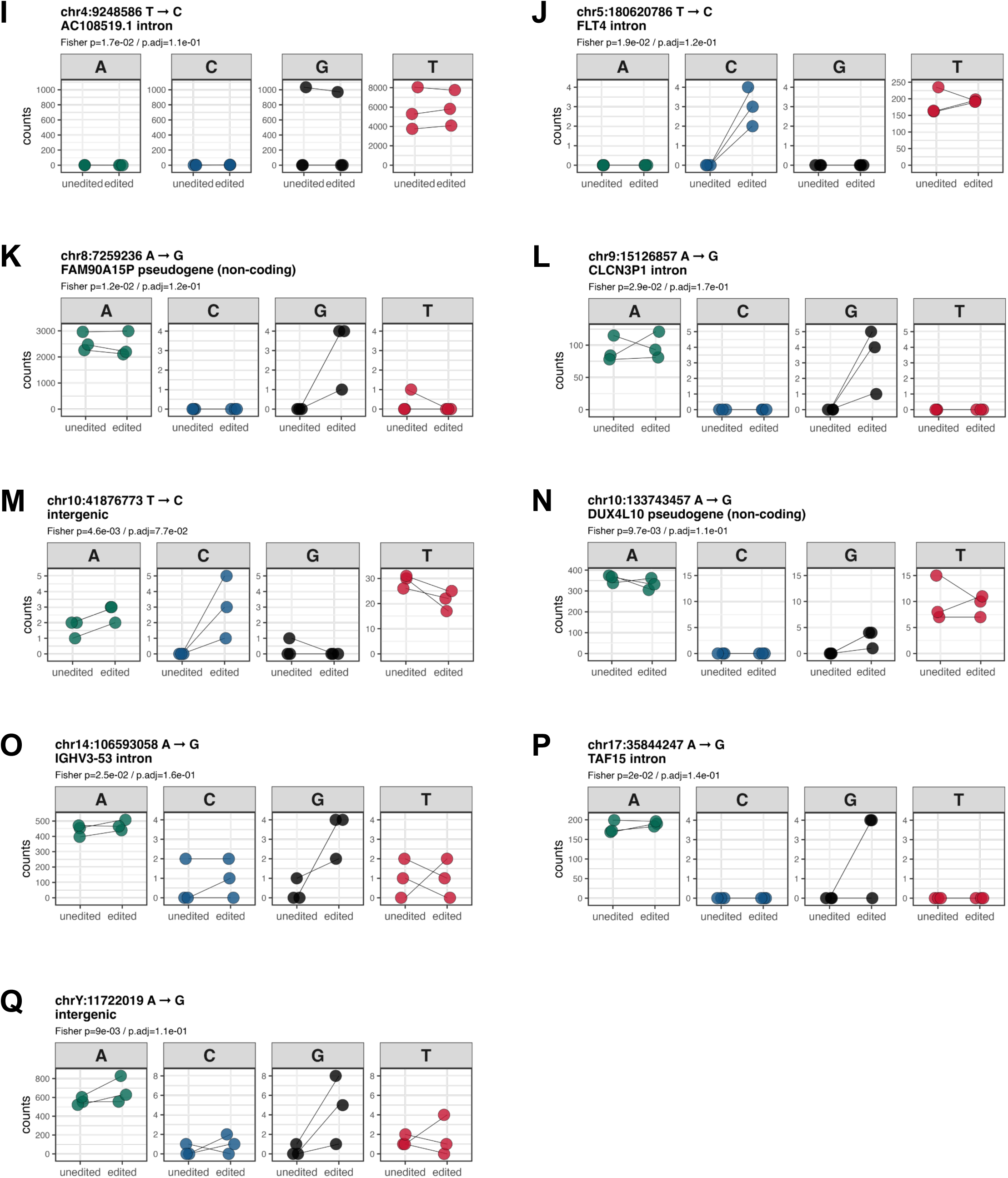

## References

1. Rocha, V., Wagner, J.E., Sobocinski, K.A., Klein, J.P., Zhang, M.J., Horowitz, M.M., and Gluckman, E. (2000). Graft-versus-host disease in children who have received a cord-blood or bone marrow transplant from an HLA-identical sibling. Eurocord and International Bone Marrow Transplant Registry Working Committee on Alternative Donor and Stem Cell Sources. N Engl J Med 342, 1846–1854. 10.1056/NEJM200006223422501.

2. Verneris, M.R., Brunstein, C.G., Barker, J., MacMillan, M.L., DeFor, T., McKenna, D.H., Burke, M.J., Blazar, B.R., Miller, J.S., McGlave, P.B., et al. (2009). Relapse risk after umbilical cord blood transplantation: enhanced graft-versus-leukemia effect in recipients of 2 units. Blood 114, 4293–4299. 10.1182/blood-2009-05-220525.

3. Horgan, C., Mullanfiroze, K., Rauthan, A., Patrick, K., Butt, N.A., Mirci-Danicar, O., O’Connor, O., Furness, C., Deshpande, A., Lawson, S., et al. (2023). T-cell replete cord transplants give superior outcomes in high-risk and relapsed/refractory pediatric myeloid malignancy. Blood Adv 7, 2155–2165. 10.1182/bloodadvances.2022009253.

4. Ballen, K.K., Gluckman, E., and Broxmeyer, H.E. (2013). Umbilical cord blood transplantation: the first 25 years and beyond. Blood 122, 491–498. 10.1182/blood-2013-02-453175.

5. Fares, I., Chagraoui, J., Gareau, Y., Gingras, S., Ruel, R., Mayotte, N., Csaszar, E., Knapp, D.J.H.F., Miller, P., Ngom, M., et al. (2014). Cord blood expansion. Pyrimidoindole derivatives are agonists of human hematopoietic stem cell self-renewal. Science 345, 1509– 1512. 10.1126/science.1256337.

6. Chagraoui, J., Girard, S., Spinella, J.-F., Simon, L., Bonneil, E., Mayotte, N., MacRae, T., Coulombe-Huntington, J., Bertomeu, T., Moison, C., et al. (2021). UM171 Preserves Epigenetic Marks that Are Reduced in Ex Vivo Culture of Human HSCs via Potentiation of the CLR3-KBTBD4 Complex. Cell Stem Cell 28, 48–62.e6. 10.1016/j.stem.2020.12.002.

7. Chagraoui, J., Girard, S., Mallinger, L., Mayotte, N., Tellechea, M.F., and Sauvageau, G. (2024). KBTBD4-mediated reduction of MYC is critical for hematopoietic stem cell expansion upon UM171 treatment. Blood 143, 882–894. 10.1182/blood.2023021342.

8. Cohen, S., Roy, J., Lachance, S., Delisle, J.-S., Marinier, A., Busque, L., Roy, D.-C., Barabé, F., Ahmad, I., Bambace, N., et al. (2020). Hematopoietic stem cell transplantation using single UM171-expanded cord blood: a single-arm, phase 1-2 safety and feasibility study. Lancet Haematol 7, e134–e145. 10.1016/S2352-3026(19)30202-9.

9. Dumont-Lagacé, M., Feghaly, A., Meunier, M.-C., Finney, M., Van’t Hof, W., Masson Frenet, E., Sauvageau, G., and Cohen, S. (2022). UM171 Expansion of Cord Blood Improves Donor Availability and HLA Matching For All Patients, Including Minorities. Transplant Cell Ther 28, 410.e1–410.e5. 10.1016/j.jtct.2022.03.016.

10. Cohen, S., Bambace, N., Ahmad, I., Roy, J., Tang, X., Zhang, M.-J., Burns, L., Barabé, F., Bernard, L., Delisle, J.-S., et al. (2023). Improved outcomes of UM171-expanded cord blood transplantation compared with other graft sources: real-world evidence. Blood Adv 7, 5717– 5726. 10.1182/bloodadvances.2023010599.

11. Dumont-Lagacé, M., Li, Q., Tanguay, M., Chagraoui, J., Kientega, T., Cardin, G.B., Brasey, A., Trofimov, A., Carli, C., Ahmad, I., et al. (2021). UM171-Expanded Cord Blood Transplants Support Robust T Cell Reconstitution with Low Rates of Severe Infections. Transplant Cell Ther 27, 76.e1–76.e9. 10.1016/j.bbmt.2020.09.031.

12. Rajvanshi, P., Shulman, H.M., Sievers, E.L., and McDonald, G.B. (2002). Hepatic sinusoidal obstruction after gemtuzumab ozogamicin (Mylotarg) therapy. Blood 99, 2310–2314. 10.1182/blood.v99.7.2310.

13. Godwin, C.D., Gale, R.P., and Walter, R.B. (2017). Gemtuzumab ozogamicin in acute myeloid leukemia. Leukemia 31, 1855–1868. 10.1038/leu.2017.187.

14. Krupka, C., Kufer, P., Kischel, R., Zugmaier, G., Bögeholz, J., Köhnke, T., Lichtenegger, F.S., Schneider, S., Metzeler, K.H., Fiegl, M., et al. (2014). CD33 target validation and sustained depletion of AML blasts in long-term cultures by the bispecific T-cell-engaging antibody AMG 330. Blood 123, 356–365. 10.1182/blood-2013-08-523548.

15. Ravandi, F., Subklewe, M., Walter, R.B., Vachhani, P., Ossenkoppele, G., Buecklein, V., Döhner, H., Jongen-Lavrencic, M., Baldus, C.D., Fransecky, L., et al. (2024). Safety and tolerability of AMG 330 in adults with relapsed/refractory AML: a phase 1a dose-escalation study. Leuk Lymphoma 65, 1281–1291. 10.1080/10428194.2024.2346755.

16. Tambaro, F.P., Singh, H., Jones, E., Rytting, M., Mahadeo, K.M., Thompson, P., Daver, N., DiNardo, C., Kadia, T., Garcia-Manero, G., et al. (2021). Autologous CD33-CAR-T cells for treatment of relapsed/refractory acute myelogenous leukemia. Leukemia 35, 3282–3286. 10.1038/s41375-021-01232-2.

17. Haubner, S., Subklewe, M., and Sadelain, M. (2025). Honing CAR T cells to tackle acute myeloid leukemia. Blood 145, 1113–1125. 10.1182/blood.2024024063.

18. Kim, M.Y., Yu, K.-R., Kenderian, S.S., Ruella, M., Chen, S., Shin, T.-H., Aljanahi, A.A., Schreeder, D., Klichinsky, M., Shestova, O., et al. (2018). Genetic Inactivation of CD33 in Hematopoietic Stem Cells to Enable CAR T Cell Immunotherapy for Acute Myeloid Leukemia. Cell 173, 1439–1453.e19.

19. Borot, F., Wang, H., Ma, Y., Jafarov, T., Raza, A., Ali, A.M., and Mukherjee, S. (2019). Gene-edited stem cells enable CD33-directed immune therapy for myeloid malignancies. Proc Natl Acad Sci U S A 116, 11978–11987. 10.1073/pnas.1819992116.

20. Petty, N.E., Radtke, S., Kanestrom, G., Fields, E., Humbert, O., Fiorenza, S., Llewellyn, M.J., Laszlo, G.S., Thomas, J., Burger, Z., et al. (2025). Protection of CD33-modified hematopoietic stem cell progeny from CD33-directed CAR T cells in nonhuman primates. Blood Adv, bloodadvances.2024015016. 10.1182/bloodadvances.2024015016.

21. Humbert, O., Laszlo, G.S., Sichel, S., Ironside, C., Haworth, K.G., Bates, O.M., Beddoe, M.E., Carrillo, R.R., Kiem, H.-P., and Walter, R.B. (2019). Engineering resistance to CD33-targeted immunotherapy in normal hematopoiesis by CRISPR/Cas9-deletion of CD33 exon 2. Leukemia 33, 762–808. 10.1038/s41375-018-0277-8.

22. 22. Brinkman-Van der Linden, E.C.M., Angata, T., Reynolds, S.A., Powell, L.D., Hedrick, S.M., and Varki, A. (2003). CD33/Siglec-3 binding specificity, expression pattern, and consequences of gene deletion in mice. Mol Cell Biol 23, 4199–4206. 10.1128/MCB.23.12.4199-4206.2003.

23. Lydeard, J.R., Lin, M.I., Ge, H.G., Halfond, A., Wang, S., Jones, M.B., Etchin, J., Angelini, G., Xavier-Ferrucio, J., Lisle, J., et al. (2023). Development of a gene edited next-generation hematopoietic cell transplant to enable acute myeloid leukemia treatment by solving off-tumor toxicity. Mol Ther Methods Clin Dev 31, 101135. 10.1016/j.omtm.2023.101135.

24. Lehnertz, B., Chagraoui, J., MacRae, T., Tomellini, E., Corneau, S., Mayotte, N., Boivin, I., Durand, A., Gracias, D., and Sauvageau, G. (2021). HLF Expression Defines the Human Hematopoietic Stem Cell State. Blood, blood.2021010745. 10.1182/blood.2021010745.

25. Billon, P., Bryant, E.E., Joseph, S.A., Nambiar, T.S., Hayward, S.B., Rothstein, R., and Ciccia, A. (2017). CRISPR-Mediated Base Editing Enables Efficient Disruption of Eukaryotic Genes through Induction of STOP Codons. Mol Cell 67, 1068–1079.e4. 10.1016/j.molcel.2017.08.008.

26. Komor, A.C., Kim, Y.B., Packer, M.S., Zuris, J.A., and Liu, D.R. (2016). Programmable editing of a target base in genomic DNA without double-stranded DNA cleavage. Nature 533, 420–424. 10.1038/nature17946.

27. Crocker, P.R., Paulson, J.C., and Varki, A. (2007). Siglecs and their roles in the immune system. Nat Rev Immunol 7, 255–266. 10.1038/nri2056.

28. Casirati, G., Cosentino, A., Mucci, A., Salah Mahmoud, M., Ugarte Zabala, I., Zeng, J., Ficarro, S.B., Klatt, D., Brendel, C., Rambaldi, A., et al. (2023). Epitope editing enables targeted immunotherapy of acute myeloid leukaemia. Nature 621, 404–414. 10.1038/s41586-023-06496-5.

29. Marone, R., Landmann, E., Devaux, A., Lepore, R., Seyres, D., Zuin, J., Burgold, T., Engdahl, C., Capoferri, G., Dell’Aglio, A., et al. (2023). Epitope-engineered human hematopoietic stem cells are shielded from CD123-targeted immunotherapy. J Exp Med 220, e20231235. 10.1084/jem.20231235.

30. Wellhausen, N., O’Connell, R.P., Lesch, S., Engel, N.W., Rennels, A.K., Gonzales, D., Herbst, F., Young, R.M., Garcia, K.C., Weiner, D., et al. (2023). Epitope base editing CD45 in hematopoietic cells enables universal blood cancer immune therapy. Sci Transl Med 15, eadi1145. 10.1126/scitranslmed.adi1145.

31. Garaudé, S., Marone, R., Lepore, R., Devaux, A., Beerlage, A., Seyres, D., Dell’ Aglio, A., Juskevicius, D., Zuin, J., Burgold, T., et al. (2024). Selective haematological cancer eradication with preserved haematopoiesis. Nature 630, 728–735. 10.1038/s41586-024-07456-3.

32. Pérez-Oliva, A.B., Martínez-Esparza, M., Vicente-Fernández, J.J., Corral-San Miguel, R., García-Peñarrubia, P., and Hernández-Caselles, T. (2011). Epitope mapping, expression and post-translational modifications of two isoforms of CD33 (CD33M and CD33m) on lymphoid and myeloid human cells. Glycobiology 21, 757–770. 10.1093/glycob/cwq220.

33. Walton, R.T., Christie, K.A., Whittaker, M.N., and Kleinstiver, B.P. (2020). Unconstrained genome targeting with near-PAMless engineered CRISPR-Cas9 variants. Science 368, 290–296. 10.1126/science.aba8853.

34. Richter, M.F., Zhao, K.T., Eton, E., Lapinaite, A., Newby, G.A., Thuronyi, B.W., Wilson, C., Koblan, L.W., Zeng, J., Bauer, D.E., et al. (2020). Phage-assisted evolution of an adenine base editor with improved Cas domain compatibility and activity. Nature biotechnology 38, 883–891.

35. Neugebauer, M.E., Hsu, A., Arbab, M., Krasnow, N.A., McElroy, A.N., Pandey, S., Doman, J.L., Huang, T.P., Raguram, A., Banskota, S., et al. (2023). Evolution of an adenine base editor into a small, efficient cytosine base editor with low off-target activity. Nat Biotechnol 41, 673–685. 10.1038/s41587-022-01533-6.

36. Shaw, B.C., and Estus, S. (2021). Pseudogene-Mediated Gene Conversion After CRISPR-Cas9 Editing Demonstrated by Partial CD33 Conversion with SIGLEC22P. CRISPR J 4, 699–709. 10.1089/crispr.2021.0052.

37. Chen, L., Zhang, S., Xue, N., Hong, M., Zhang, X., Zhang, D., Yang, J., Bai, S., Huang, Y., Meng, H., et al. (2023). Engineering a precise adenine base editor with minimal bystander editing. Nat Chem Biol 19, 101–110. 10.1038/s41589-022-01163-8.

38. Perrotta, R.M., Vinke, S., Ferreira, R., Moret, M., Mahas, A., Chiappino-Pepe, A., Riedmayr, L.M., Mehra, A.-T., Lehmann, L.S., and Church, G.M. (2024). Machine Learning and Directed Evolution of Base Editing Enzymes. Preprint at Synthetic Biology, 10.1101/2024.05.17.594556.

39. Läubli, H., and Varki, A. (2020). Sialic acid-binding immunoglobulin-like lectins (Siglecs) detect self-associated molecular patterns to regulate immune responses. Cell Mol Life Sci 77, 593–605. 10.1007/s00018-019-03288-x.

40. Lin, S.-Y., Schmidt, E.N., Takahashi-Yamashiro, K., and Macauley, M.S. (2025). Roles for Siglec-glycan interactions in regulating immune cells. Semin Immunol 77, 101925. 10.1016/j.smim.2024.101925.

41. Rillahan, C.D., Macauley, M.S., Schwartz, E., He, Y., McBride, R., Arlian, B.M., Rangarajan, J., Fokin, V.V., and Paulson, J.C. (2014). Disubstituted Sialic Acid Ligands Targeting Siglecs CD33 and CD22 Associated with Myeloid Leukaemias and B Cell Lymphomas. Chem Sci 5, 2398–2406. 10.1039/C4SC00451E.

42. Bhattacherjee, A., Daskhan, G.C., Bains, A., Watson, A.E.S., Eskandari-Sedighi, G., St Laurent, C.D., Voronova, A., and Macauley, M.S. (2021). Increasing phagocytosis of microglia by targeting CD33 with liposomes displaying glycan ligands. J Control Release 338, 680–693. 10.1016/j.jconrel.2021.09.010.

43. Movsisyan, L.D., and Macauley, M.S. (2020). Structural advances of Siglecs: insight into synthetic glycan ligands for immunomodulation. Org Biomol Chem 18, 5784–5797. 10.1039/d0ob01116a.

44. Vakulskas, C.A., Dever, D.P., Rettig, G.R., Turk, R., Jacobi, A.M., Collingwood, M.A., Bode, N.M., McNeill, M.S., Yan, S., Camarena, J., et al. (2018). A high-fidelity Cas9 mutant delivered as a ribonucleoprotein complex enables efficient gene editing in human hematopoietic stem and progenitor cells. Nature Medicine 24, 1216–1224.

45. Bae, S., Park, J., and Kim, J.-S. (2014). Cas-OFFinder: a fast and versatile algorithm that searches for potential off-target sites of Cas9 RNA-guided endonucleases. Bioinformatics 30, 1473–1475. 10.1093/bioinformatics/btu048.

46. Concordet, J.-P., and Haeussler, M. (2018). CRISPOR: intuitive guide selection for CRISPR/Cas9 genome editing experiments and screens. Nucleic Acids Res 46, W242– W245. 10.1093/nar/gky354.

47. Gerstung, M., Beisel, C., Rechsteiner, M., Wild, P., Schraml, P., Moch, H., and Beerenwinkel, N. (2012). Reliable detection of subclonal single-nucleotide variants in tumour cell populations. Nat Commun 3, 811. 10.1038/ncomms1814.

48. Tsai, S.Q., Zheng, Z., Nguyen, N.T., Liebers, M., Topkar, V.V., Thapar, V., Wyvekens, N., Khayter, C., Iafrate, A.J., Le, L.P., et al. (2015). GUIDE-seq enables genome-wide profiling of off-target cleavage by CRISPR-Cas nucleases. Nat Biotechnol 33, 187–197. 10.1038/nbt.3117.

49. Baron, J., and Wang, E.S. (2018). Gemtuzumab ozogamicin for the treatment of acute myeloid leukemia. Expert Rev Clin Pharmacol 11, 549–559. 10.1080/17512433.2018.1478725.

50. Schiroli, G., Conti, A., Ferrari, S., della Volpe, L., Jacob, A., Albano, L., Beretta, S., Calabria, A., Vavassori, V., Gasparini, P., et al. (2019). Precise Gene Editing Preserves Hematopoietic Stem Cell Function following Transient p53-Mediated DNA Damage Response. Cell stem cell 24, 551–565.e8.

51. Maganti, H.B., Bailey, A.J.M., Kirkham, A.M., Shorr, R., Pineault, N., and Allan, D.S. (2021). Persistence of CRISPR/Cas9 gene edited hematopoietic stem cells following transplantation: A systematic review and meta-analysis of preclinical studies. Stem Cells Transl Med 10, 996–1007. 10.1002/sctm.20-0520.

52. Ferrari, S., Vavassori, V., Canarutto, D., Jacob, A., Castiello, M.C., Javed, A.O., and Genovese, P. (2021). Gene Editing of Hematopoietic Stem Cells: Hopes and Hurdles Toward Clinical Translation. Front Genome Ed 3, 618378. 10.3389/fgeed.2021.618378.

53. Lee, B.-C., Lozano, R.J., and Dunbar, C.E. (2021). Understanding and overcoming adverse consequences of genome editing on hematopoietic stem and progenitor cells. Mol Ther 29, 3205–3218. 10.1016/j.ymthe.2021.09.001.

54. Knapp, D.J.H.F., Hammond, C.A., Hui, T., Loenhout, M.T.J., Wang, F., Aghaeepour, N., Miller, P.H., Moksa, M., Rabu, G.M., Beer, P.A., et al. (2018). Single-cell analysis identifies a CD33 + subset of human cord blood cells with high regenerative potential. Nature Cell Biology 20, 710–720.

55. Li, W., Köster, J., Xu, H., Chen, C.-H., Xiao, T., Liu, J.S., Brown, M., and Liu, X.S. (2015). Quality control, modeling, and visualization of CRISPR screens with MAGeCK-VISPR. Genome Biol 16, 281. 10.1186/s13059-015-0843-6.

56. Langmead, B., Trapnell, C., Pop, M., and Salzberg, S.L. (2009). Ultrafast and memory-efficient alignment of short DNA sequences to the human genome. Genome Biol 10, R25. 10.1186/gb-2009-10-3-r25.

